# Dysregulation of murine long non-coding single cell transcriptome in non-alcoholic steatohepatitis and liver fibrosis

**DOI:** 10.1101/2022.12.26.521944

**Authors:** Kritika Karri, David J. Waxman

## Abstract

LncRNAs are typified by low expression, nuclear enrichment, high tissue-specificity and functional diversity, but the vast majority are uncharacterized. Here, we develop a catalog of 48,000 mouse liver-expressed lncRNAs and use to elucidate lncRNA dysregulation in high fat diet-induced non-alcoholic steatohepatitis and carbon tetrachloride-induced liver fibrosis by single cell RNA-seq. LncRNA zonation patterns across the liver lobule were discovered in five cell populations. Perturbations in lncRNA expression were common in disease-associated cell types, including non-alcoholic steatohepatitis-associated macrophages, a hallmark of fatty liver disease progression, and collagen-producing myofibroblasts, key to liver fibrosis. Gene regulatory network analysis linked individual lncRNAs to biological pathways and network centrality metrics identified network-essential, disease-associated regulatory lncRNAs. Regulatory lncRNAs with network-predicted target gene promoters enriched for triplex-based lncRNA binding were also identified. These findings elucidate hepatic lncRNA cell-type specificities, spatial zonation patterns, linked biological networks and dysregulation in disease progression. A subset of the disease-associated lncRNAs have human orthologs and are promising candidates for biomarkers and therapeutic targets.

## Introduction

Nonalcoholic fatty liver disease (NAFLD) is rapidly becoming the most common chronic liver disease, affecting 25% of the world’s adult population, most notably individuals with obesity, type II diabetes and metabolic syndrome^1^. NAFLD comprises a continuum of liver pathologies, ranging from fat accumulation, known as simple steatosis or nonalcoholic fatty liver, to non-alcoholic steatohepatitis (NASH), a more severe disease subtype characterized by excessive lipid accumulation, chronic inflammation, hepatocyte ballooning and varying degrees of fibrosis^2, 3^. Liver fibrosis is characterized by excessive accumulation of collagens and other extracellular matrix proteins^4^ and can be induced in NASH or by environmental chemical-induced injury and alcohol abuse^5^. The underlying mechanisms of development and progression of these liver diseases are still poorly understood. Moreover, there are no approved therapeutics for advanced NASH and liver fibrosis^6^, which all too frequently advance to liver cirrhosis and hepatocellular carcinoma^7, 8^.

Long non-coding RNAs (lncRNAs) comprise a heterogeneous class of RNA-encoding genes, primarily defined by their low protein coding potential/low translational activity and by a minimum RNA length of 200 nt. LncRNAs often have low expression, show strong nuclear enrichment^9^ have high tissue-specificity^10^, and may regulate gene expression through effects on chromatin states and transcriptional regulation or at the post-transcriptional level^11, 12^. Thousands of liver-expressed lncRNAs have been identified, a subset of which are responsive to endogenous hormones^13–15^ or exposure to xenobiotics^16–19^, many of which can promote NASH, cirrhosis and other liver pathologies^20–22^. Prior studies identified individual lncRNAs that impact liver disease^23–25^; examples include SRA, which promotes hepatic steatosis by repressing adipose triglyceride lipase expression^26^, GAS5, which attenuates carbon tetrachloride (CCl_4_)-induced liver fibrosis by acting as a sponge for miRNA-23a^27^, and HULC, which inhibits liver fibrosis associated with NAFLD^28^. Thousands of other liver-expressed lncRNAs are uncharacterized or even unidentified, many of which are likely to impact liver pathophysiology.

Hepatocytes account for 60-70% of all cells in the liver, with the balance largely comprised of three major non- parenchymal cell types: endothelial cells, hepatic stellate cells, and Kupffer cells (liver resident macrophages). Liver cell type-specific gene expression patterns and their zonated regulation across the liver lobule have been elucidated in both healthy liver^29–31^ and in high fat diet-induced liver disease^32–34^ by using single cell (sc)RNA- sequencing technologies. scRNA-seq has also elucidated the role of hepatic mesenchymal cells, including hepatic^35–38^ stellate cells (HSCs), in liver fibrosis induced by hepatotoxins such as CCl_4_ (Fig. 1A). While these studies have determined the roles of liver cell subpopulations and individual protein coding genes (PCGs)^39^, prior studies of lncRNAs have largely been limited to IncRNAs with RefSeq gene or other annotations, which comprise only a small fraction of the tens of thousands of lncRNAs thought to be encoded by the genome. Many lncRNAs are expressed in bulk tissue at much lower levels than PCGs, but often exhibit high tissue specificity, raising the possibility that detection sensitivity may actually be increased by using scRNA-seq technology to characterize lncRNAs whose expression is restricted to a specific subpopulation of cells in the liver.

**Fig. 1.**
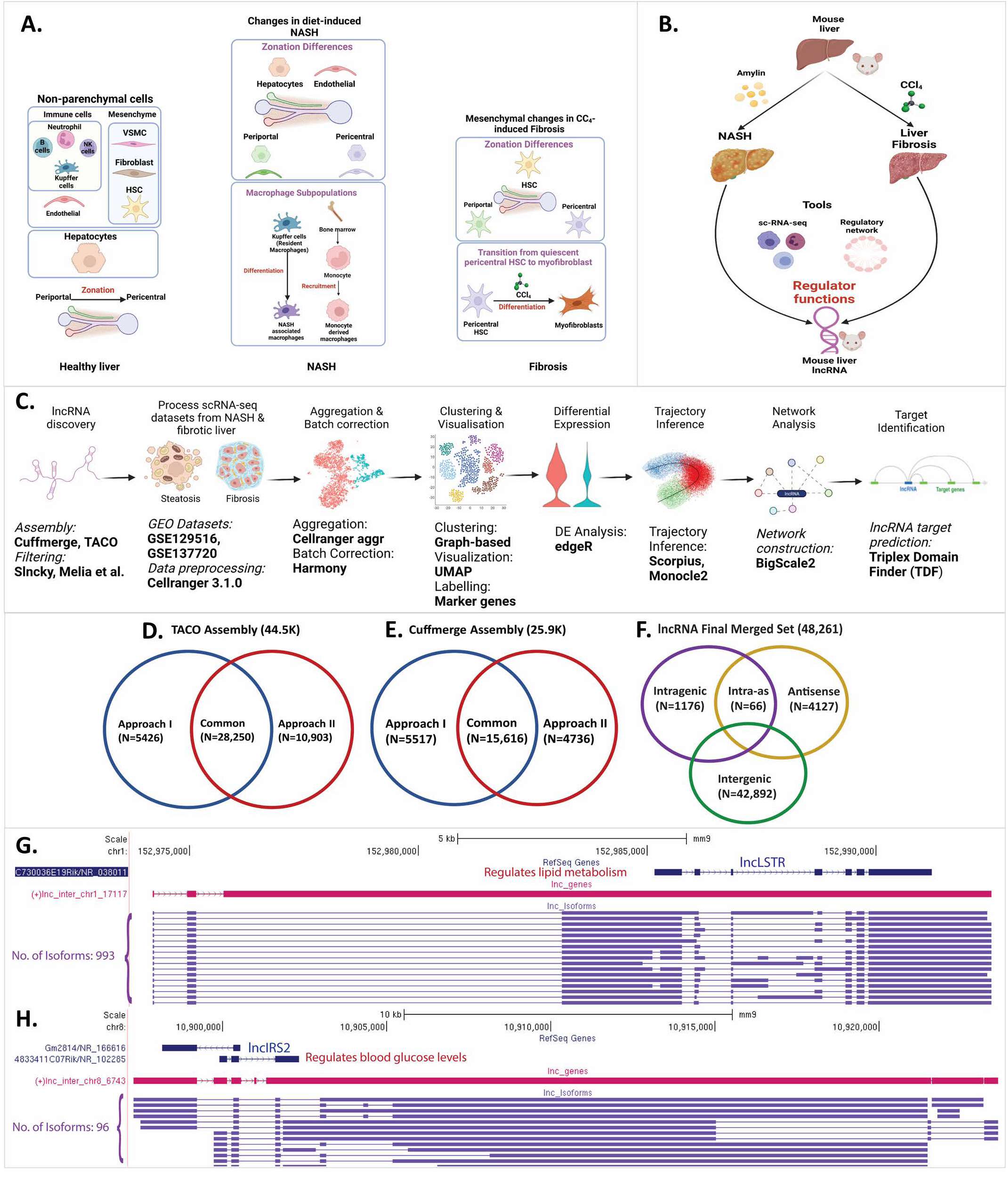
LncRNA discovery and role in NASH and liver fibrosis analyzed using single cell technology. **A.** Healthy liver is comprised of hepatocytes and non-parenchymal cells, notably endothelial cells, mesenchymal cells, and immune cell populations. With the emergence of NASH, changes in gene expression and zonation occur in hepatocytes and endothelial cells and new macrophage subpopulations emerge. In CCl_4_-induced liver fibrosis, changes in hepatic mesenchymal cells include zonation differences in hepatic stellate cells and the transition of pericentral stellate cells to collagen-producing myofibroblasts. **B.** Investigation of regulatory roles of lncRNAs in two liver disease models: diet-induce NASH and CCl_4_-induced liver fibrosis. **C.** Computational workflow for characterization of functional roles of liver-expressed lncRNAs used in this study. **D, E.** Numbers of mouse (mm9) liver lncRNAs discovered using TACO assembly (D) and Cuffmerge assembly (E), with two different filtering approaches, I and II. **F.** Final set of 48,261 mouse lncRNAs, classified based on their location with respect to PCGs after conversion to mouse mm10 genomic coordinates. **G, H.** Two liver-expressed lncRNAs, both found to be comprised of many novel isoforms, a subset of which is shown.

Here, we sought to elucidate on a global scale the roles of liver-expressed lncRNAs in biological pathways related to liver disease development. We used an integrative approach to assemble the hepatic transcriptome from >2,000 bulk murine liver RNA-seq samples and discovered more than 48,000 liver-expressed IncRNAs, a majority novel and previously uncharacterized, including many lncRNAs that share orthology with corresponding human sequences. We integrated multiple public scRNA-seq datasets for healthy mouse liver to create a single-cell transcriptomic reference atlas, which enabled us to characterize the liver cell type specificities of thousands of novel lncRNAs and identify more than 100 liver cell type-specific IncRNA marker genes. We elucidated liver lncRNA zonation profiles using trajectory inference algorithms for 5 liver cell populations, complementing those reported earlier for PCGs^35, 40–42^ (Fig. 1A). Further, to elucidate lncRNA dysregulation in liver disease, we analyzed liver scRNA-seq data from mice fed a high fat, high fructose diet (HFHFD, also known as AMLN diet)^33^, which induces disease progression from healthy liver to NAFLD (simple steatosis) and then NASH, revealing IncRNA transcriptomic dysregulation patterns during disease progression. We also analyzed the role of lncRNAs in the hepatic mesenchyme from healthy and CCl_4_-induced fibrotic liver^35^ (Fig. 1B). Finally, we constructed gene regulatory networks for both healthy and diseased liver to discover key regulatory lncRNAs based on gene network centrality metrics, and we identified a subset of regulatory lncRNAs whose PCG target gene promoters are significantly enriched for direct interactions via triplex-based lncRNA binding (Fig. 1C). Overall, our findings reveal an unanticipated complexity of hepatic IncRNA biology. The datasets obtained are expected to serve as a rich resource for discovery of lncRNA biomarkers and in studies targeting IncRNAs implicated in development of NASH and foreign chemical-induced liver fibrosis.

## Results

### Global discovery of liver-expressed lncRNAs

We reconstructed the mouse liver transcriptome from 2,089 bulk RNA-seq samples representing a wide range of biological conditions (Table S1A). We integrated results from two different transcriptome assembly methods, TACO^43^ and Cuffmerge^44^, and used two approaches to analyze the output and identify liver-expressed lncRNA genes and isoform structures^14, 45^. The transcriptomes generated by each method were each processed and analyzed using two discovery pipelines (Fig. 1D, Fig. 1E), which after integrating with a prior set of 15,558 mouse liver lncRNAs based on a much smaller number of bulk RNA-seq samples^13^ yielded a global set of 48,261 liver-expressed lncRNAs, including 5,656 multi-exonic genes (Table S1C) and a total of 150,280 isoforms (Table S1D). 89% of the 48,261 IncRNAs are intergenic (Fig. 1F). Further, 9,543 (19.8%) of the lncRNAs have orthologous sequences in the human genome (hg38), and 1,722 were orthologous to an incomplete set comprised of 5,795 lncRNAs that we previously identified in rat liver^18^ (Table S1E). The final lncRNA dataset includes many novel isoforms for some well-characterized liver lncRNAs. For example, we identified hundreds of isoforms of LncLSTR (lnc17117), a liver-enriched RNA that regulates lipid metabolism in mice^46^ (Fig. 1G) and 96 isoforms of LincIRS2 (lnc6743), which protects against diabetes and whose knockdown increases blood glucose, insulin resistance and aberrant glucose output^47^ (Fig. 1H). However, the vast majority of the 48,261 liver lncRNAs are novel and of unknown function.

### LncRNA expression in liver cell subpopulations

The low expression of many lncRNAs makes it difficult to reliably characterize their expression using bulk tissue RNA-seq, where expression patterns are dominated by hepatocytes, which comprise 60-70% of all cells in adult liver^48, 49^. Moreover, the high frequency of scRNA-seq drop out of low abundance transcripts limits the ability of scRNA-seq to detect and characterize the many thousands of low expressed lncRNAs. To increase the sensitivity for IncRNA detection, we pooled, integrated, and harmonized data from four public mouse liver scRNA-seq datasets comprising 39,878 liver cells (Table S1G) to give a single, uniform single-cell landscape for healthy mouse liver. The resulting UMAP, which is enriched in non-parenchymal cells (Fig. 2A), contains 13 major cell clusters identified by their marker gene expression patterns (Fig. S1A): hepatocytes, endothelial cells, Kupffer cells (liver macrophages), mesenchymal subpopulations comprised of hepatic stellate cells (HSCs), vascular smooth muscle cells (VSMCs) and fibroblasts, as well as dendritic cells, cholangiocytes, natural killer and T cells, B cells, B plasma cells, neutrophils, and dividing cells. A total of 30,092 distinct IncRNAs were detectable in this healthy liver dataset, of which 24,961 (83%) are novel genes (Table S2B). 1,352 lncRNAs were expressed in at least 5% of cells from at least one cell cluster (Fig. 2B, Table S2A, Table S2B). Fourteen lncRNAs (11 with human orthologs, indicated by *) were expressed across all liver cell types at >10% cells/cluster (Table S2C). Examples of these widely expressed liver lncRNAs with established roles in liver biology and pathophysiology include: Malat1 (lnc31752*) and Norad/LINC00657 (lnc1906*), which promote hepatocellular carcinoma^50, 51^, as do Snhg8 (lnc2803*)^52^ and Dleu2 (lnc25736*)^53^; Gas5 (lnc733*), which alleviates collagen accumulation in fibrotic liver^54^; Neat1 (lnc14746*), which promotes NAFLD by facilitating hepatic lipid accumulation^55^ and promotes liver fibrosis^25^; Cyrano/Oip5os1 (lnc34166*), an essential developmental lncRNA^56^; and Pint (lnc4993), which interacts with the Polycomb repressive complex 2 and is required to target specific genes for histone-H3 K27 tri-methylation and gene repression^57^.

**Fig. 2.**
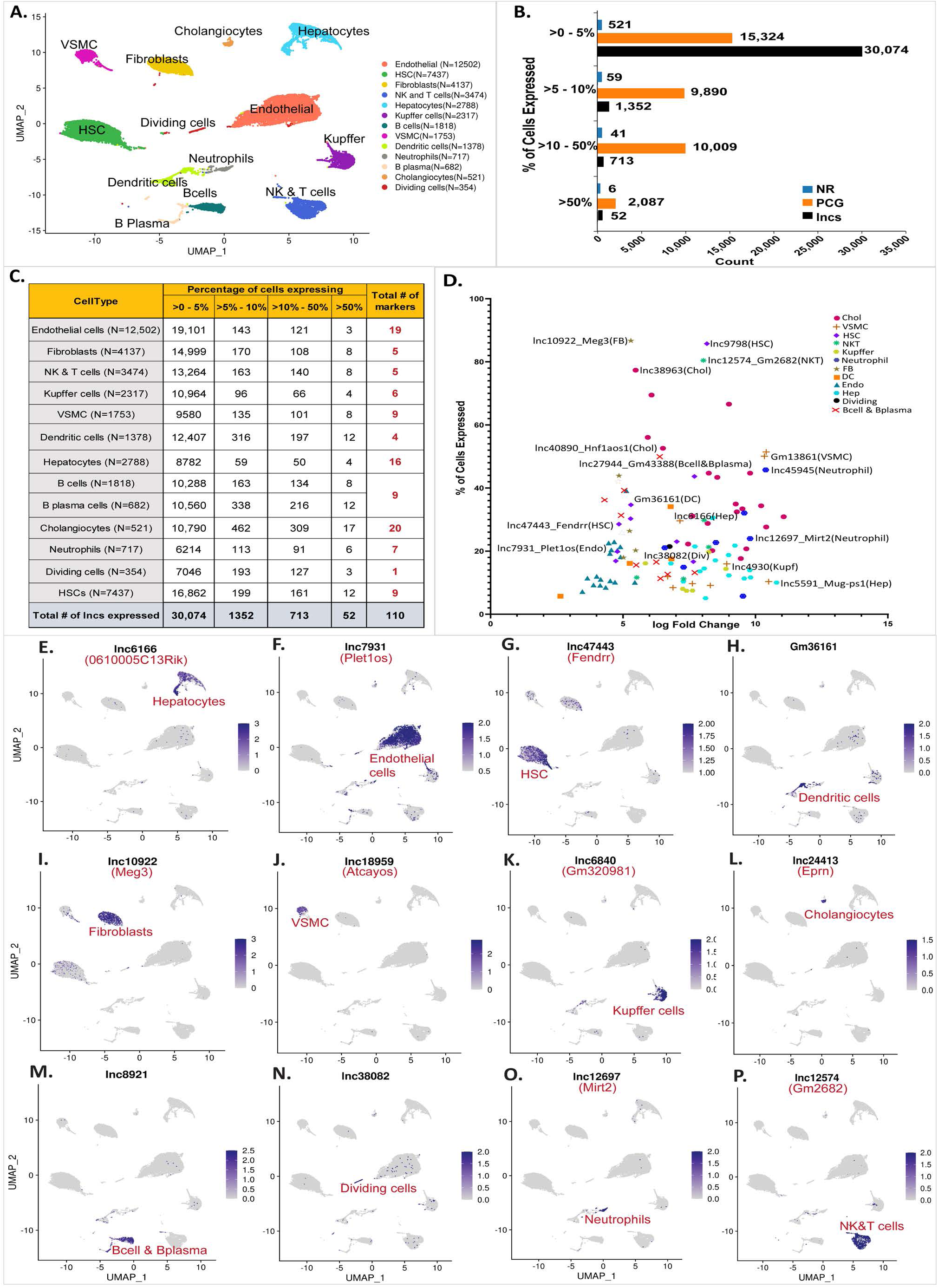
LncRNA detection in hepatic subpopulations and cell-type specific markers. **A.** UMAP of liver cell clusters based on 39,878 single-cell transcriptomes integrated from four scRNA-seq datasets. Cell counts for each cell type are indicated in parentheses and in Table S1H. **B.** Bar plots indicating number of genes in the indicated gene classes (lncRNAs, PCGs, other non-coding RNAs (NR)) that are detectably expressed in one or more liver cell types in up to 5% of cells in a cluster, in >5-10% of cells in a cluster, in >10-50% of cells in a cluster, or in >50% of cells in a cluster, based on scRNA-seq data from healthy (control) adult male mouse liver. **C.** Number of lncRNAs detected in each liver cell type, presented as a percentage of cells that express the lncRNA at ≥ 1 UMI/cell. Thus, 19,101 lncRNAs were detectably expressed in up to 5% of endothelial cells, and 3 lncRNAs were expressed in >50% of endothelial cells. Marker genes for each cell cluster (last column) are based on Table S2D. **D.** Plot depicting 110 liver cell type-specific marker lncRNAs, from panel C. X-axis, log_2_ fold-change value for differential expression of the cluster marker gene lncRNA compared to its expression in other clusters (see Methods); Y-axis, percentage of cells that express the lncRNA marker (see Table S2D). **(E-P)** Feature plots with examples of lncRNA markers from each liver cell cluster. Shown are UMAPs as in A, with color intensity indicating lncRNA expression level in individual cells.

### Cell-type specific lncRNA marker genes

110 IncRNAs showed high specificity for expression in a single liver cell type; only 44 of these lncRNAs were previously known (i.e., have RefSeq or Ensembl annotations; Table S2D, Fig. 2E-2P; see Methods),). One example is Fendrr (lnc47443*), a strong marker for HSCs (Fig. 2G). Fendrr inhibits pulmonary fibrosis^58^ and its overexpression inhibits hepatocellular carcinoma growth^59^. Another example, Meg3 (lnc10922*), has liver anti-fibrotic activity^25^ and is a marker for liver fibroblasts (Fig. 2I). Atcayos (lnc18959) is a marker for liver vascular smooth muscle cells (Fig. 2J) that regulates myogenic differentiation of satellite cells during skeletal muscle development^60^, and Ephemeron (Eprn; lnc24413) fine tunes the dynamics of the cell state transition in mouse embryonic stem cells^61^ and in liver is a cholangiocyte marker (Fig. 2L). Finally, Mirt2 (lnc12697) is a neutrophil marker (Fig. 2O) that negatively regulates inflammation^62^.

### LncRNA zonation across the healthy liver lobule

The liver is divided into small functional units called lobules fed by the hepatic artery and the portal vein, which drain to the central vein via sinusoidal capillaries. Gradients of oxygen, nutrients and hormones are established across the lobule, leading to spatial zonation of liver function and gene expression, as is seen in hepatocytes^63^, endothelial cells^40^ and HSCs^35^. Here, we elucidated the zonation patterns for both lncRNAs and PCGs in healthy liver (chow diet-fed mice) in hepatocytes and in four major non-parenchymal cell populations.

#### LncRNA zonation in hepatocytes

We used established spatial zonation markers^30^ for periportal, mid-lobular and pericentral hepatocytes in combination with trajectory inference analysis using Monocle2^64, 65^ to identify 111 IncRNAs showing significant zonation (q-val <0.001) across the liver lobule (Fig. 3A, Table S3A). For example, lnc- LFAR1, a liver-enriched lncRNA that promotes liver fibrosis by activating TGFbeta and notch signaling^25, 66^, was preferentially expressed in periportal hepatocytes, while lnc13605 and lnc14942 showed mid-lobular zonation, and Gas5 (lnc733*) and lnc6236* exemplified pericentrally zonated lncRNAs (Fig. 3A). PCGs expressed in periportal hepatocytes showed functional enrichment for specialized liver functions, such as lipid and steroid metabolism, acute inflammatory response and complement cascades, whereas pericentral hepatocytes were enriched for functions related to carboxylic acid metabolic process, oxidoreductase, peroxisome and response to xenobiotic stimulus (Table S3B), consistent with prior work^30^.

**Fig. 3.**
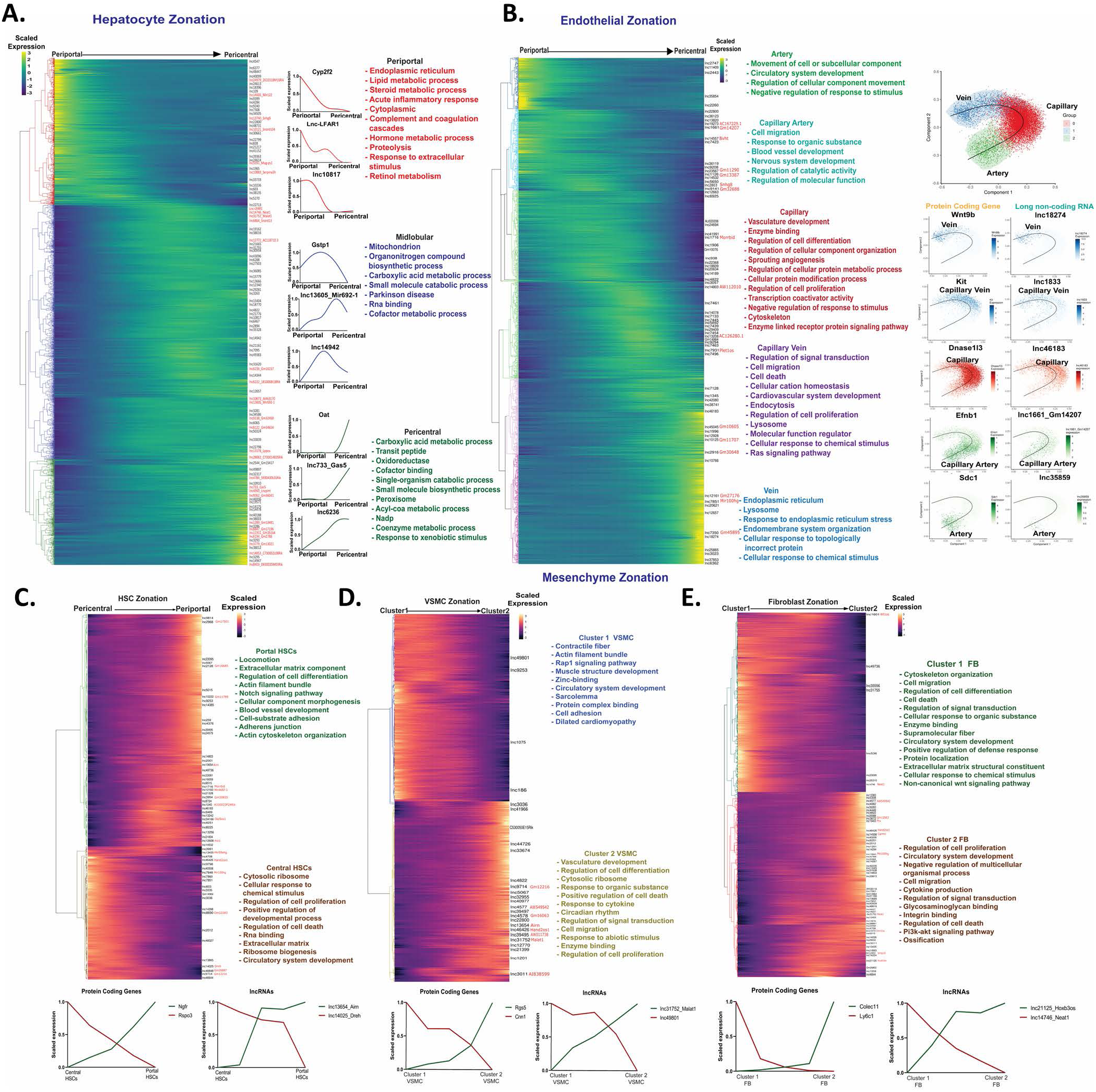
LncRNA zonation across healthy liver lobule. **A.** Heatmap showing relative expression of PCGs and lncRNAs that are zonated in hepatocytes, ordered from periportal (left) to pericentral (right), with each row corresponding to one gene, as marked at the right for lncRNAs. Right of heatmap: Zonation profiles for select genes, along with top enriched terms for each of the 3 main hepatocyte zones. Font color for enriched terms matches the dendrogram color at the left of heatmap. **B.** Heatmap showing zonated expression profiles for PCGs and lncRNAs that are zonated in endothelial cells, with clusters matching gene trajectories sequentially, from artery to capillary artery, capillary, capillary vein, and finally vein, with cluster assignments based on known spatial marker genes (see text). Top functional enrichment terms of each cluster (colored) are at the right, followed by Scorpius pseudo-time trajectories showing endothelial cell phenotypes along the artery-capillary-vein axis, with trajectory plots for select marker genes and mouse liver lncRNAs displayed across the pseudotime trajectory. **C-E.** Heatmaps showing expression trajectories for PCGs and lncRNAs that are zonated in the hepatic mesenchyme, which is comprised of HSCs (**C**), VSMCs (**D**), and fibroblasts (**E**), with top enriched terms and select marker genes shown.

#### LncRNA zonation in endothelial cells

Five zonated endothelial cell clusters were obtained and identified using established spatial marker genes for arteries (marker gene: Sdc1), capillary arteries (arterioles; Efnb1), capillaries (Dnase1I3), capillary veins (venules; Kit) and veins (Wnt9b)^42^. Overall, we identified 1,163 zonated endothelial cell transcripts, including 71 zonated lncRNAs (q-val <0.001) (Table S3C), and determined the enriched biological functions for the genes expressed in each cell cluster (Fig. 3B, Table S3D). For example, capillary endothelial cells were enriched for vasculature development and sprouting angiogenesis; while capillary vein endothelial cells were enriched for cell migration and cell death gene expression, and for Ras signaling genes, which help regulate hepatocyte zonation^67^. LncRNAs preferentially expressed in capillary arteries of the endothelium include Bvht (lnc14557), an epigenetic regulator of cardiovascular lineage commitment^68^, and Snhg8 (lnc2803*), which promotes tumorigenesis and predicts tumor recurrence in hepatocellular carcinoma^52^, while the venous cell- enriched lnc-Mir100hg (lnc7851*) activates Wnt signaling via its embedded miRNAs^69^. Other top zonation markers across the endothelium trajectory, derived using SCORPIUS^70^, are shown in Fig. 3B (right).

#### lncRNA zonation in mesenchymal cells

We investigated gene expression zonation patterns in three mesenchymal cell populations (Fig. 3C-3E): HSCs, fibroblasts, which are found in the mesenchyme surrounding the bile duct^71^, and VSMCs, which comprise a cell layer beneath endothelial cells lining the blood vessel^72^. Trajectory analysis partitioned HSCs into two sub-clusters: portal vein-associated HSCs and central vein- associated HSCs whose identities we verified using the established HSC marker genes Rspo3 (pericentral) and Ngfr (periportal)^35^ (Fig. 3C). We identified 58 HSC zonated lncRNAs, including Pvt1 (lnc12608), Airn (lnc13654) and Lnc-Dreh (lnc14025) (Fig. 3C, Table S3E). Pvt1 activates HSCs under hypoxia and promotes liver fibrosis^73, 74^ and co-zonates with periportal HSC genes, as does Airn. Pathways associated with the PCGs in this HSC cluster include locomotion, extracellular matrix, Notch signaling, and blood vessel development. In contrast, lnc-Dreh, a tumor suppressor gene for hepatocellular carcinoma^75^, was preferentially expressed in central vein-associated HSCs, which were most highly enriched for cytosolic ribosome genes and for regulation of cell proliferation (Fig. 3C, Table S3F).

The spatial zonation of smooth muscle cells has been established in brain but has not been investigated for liver. Trajectory analysis identified two distinct clusters of liver VSMCs, with 381 genes (355 PCGs, 26 lncRNAs) showing significant zonation (Table S3G). VSMC cluster 1 was characterized by high expression of Rgs5, a marker for brain pericytes^76, 77^, and was enriched in functions related to contractile fiber and muscle structure development. In contrast, VSMC cluster 2 was characterized by high expression of Cnn1, a marker for arterial smooth muscle cells in mouse brain^77^, and was enriched for a distinct set of biological processes, including vasculature development, cell differentiation, cytosolic ribosome, cell death, and response to cytokine (Fig. 3D, Table S3H).

Fibroblast heterogeneity associated with discrete anatomical positions is seen in multiple tissues^78^ but has not been characterized for liver. Trajectory inference identified two distinct fibroblast clusters in mouse liver, which may correspond to distinct, zonated cell populations, and encompassed 1,335 PCGs and 69 lncRNAs (Table S3I). Cluster 1 liver fibroblasts were enriched for cytoskeleton organization, cell migration, cell differentiation, and cell death, while cluster 2 fibroblasts were enriched for cell proliferation, circulatory system development, and extracellular matrix (Fig. 3E, Table S3J). Neat1 (lnc14746*), which promotes liver fibrosis^25^, was enriched in cluster 1 fibroblasts, while cluster 2 fibroblasts showed enrichment for Dnm3os (lnc17273*), which promotes hepatocellular carcinoma via an epigenetic mechanism^79^, and for Carmn/Mir143HG (lnc14558*), whose loss in human VSMCs is associated with enhanced proliferation and migration^80^.

### LncRNAs dysregulation in diet-induced NASH

To elucidate the role of lncRNAs in NASH progression and the development of pathogenic cell states, we analyzed scRNA-seq data^33^ for a total of 74,718 liver cells isolated from healthy mice (chow diet) and from mice fed HFHFD (high fat high fructose diet) for either 15 wk, to induce simple steatosis (NAFLD), or for 30-34 wk, to induce NASH (Table S1H). After 15 wk high fat diet (NAFLD livers), 1,477 genes, including 308 lncRNAs, were differentially expressed (>2-fold-change at FDR <0.05) in one or more of the 12 major cell clusters identified (Fig. 4A, Table S4A). These genes showed strong functional enrichment for inflammatory response, lipid metabolic process, leukocyte migration, cytokine production and innate immune response (Table S4B). After 30 wk (NASH livers), the number of differentially expressed genes increased to 2,216, including 332 lncRNAs, and then to 3,090 genes, including 459 lncRNAs, after 34 wk (Table S4C, Table S4D). Overall, a total of 677 lncRNAs were dysregulated across the time course of NAFLD to NASH progression. The top enriched pathways for the NASH-induced and NASH-repressed genes were very similar to those seen in the NAFLD (15 wk) livers (Table S4B).

**Fig. 4.**
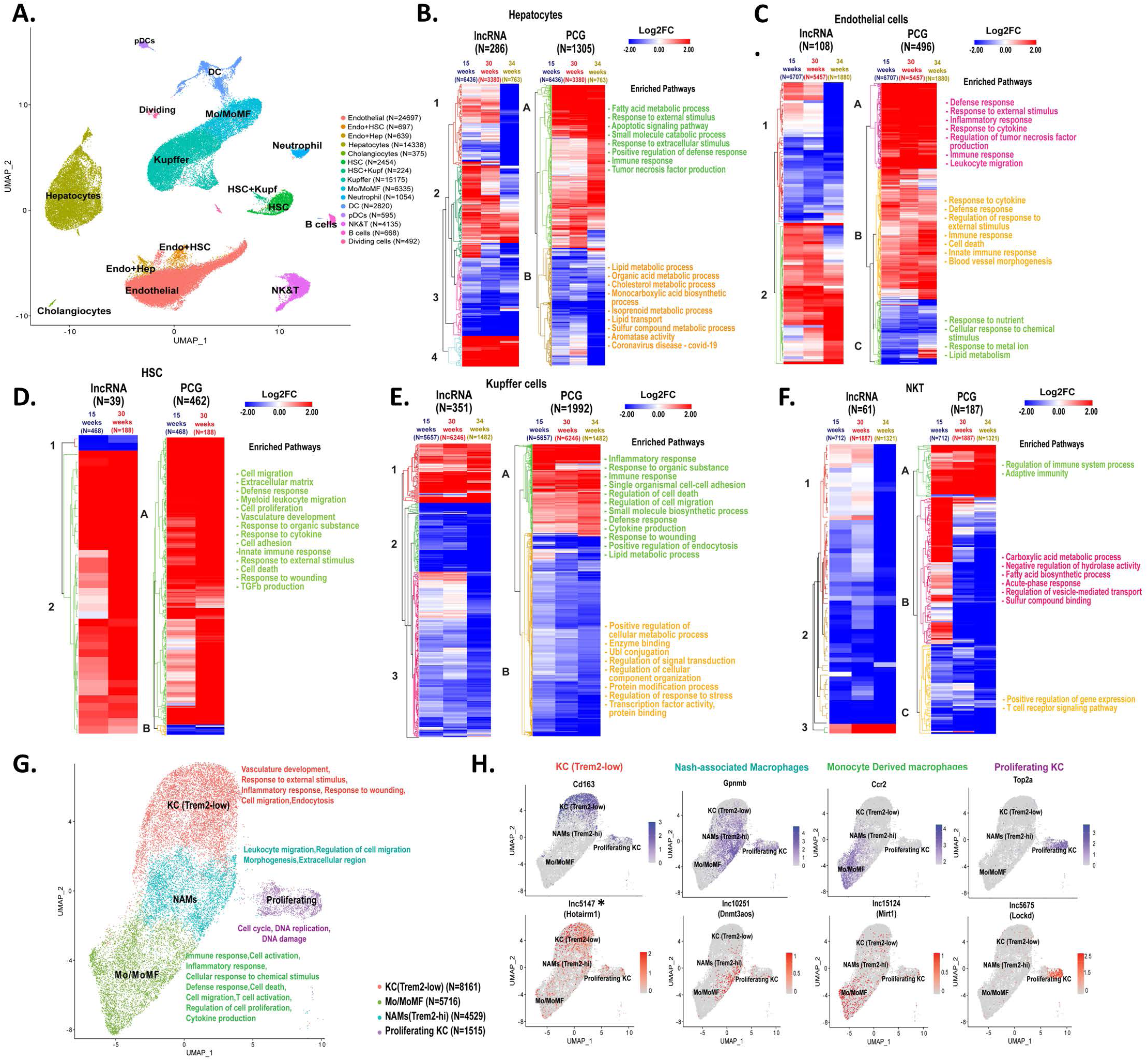
LncRNAs and PCGs perturbed across different cell types in NAFLD and NASH. **A.** UMAP showing liver cell clusters based on 74,718 cells aggregated from healthy liver (19,364 cells) with cells from livers of mice fed HFHFD for 15-wk (NAFLD liver; 23,961 cells), 30-wk (24,106 cells), or 34-wk (NASH livers; 7,287 cells) (Table S1H). Cell numbers in each cluster are shown in parenthesis. Mo/MoMF, monocytes/monocyte-derived macrophages. **(B–F)** Heat maps showing differentially expressed lncRNAs and PCGs in the indicated cells clusters from 15, 30, and 34 wk HFHFD-fed livers compared to chow diet livers, in hepatocytes **(B)**, endothelial cells **(C)**, HSCs **(D)**, Kupffer cells **(E)**, and NK & T cells **(F)**. Functional enrichment terms shown for induced and repressed PCGs in each cell type. See Table S4 for full datasets. **G.** UMAP of Kupffer (macrophage) cell subpopulations, identified based on PCG markers in each cell subtype. Kupffer cells and Mo/MoMf cell clusters shown in **A** were reclustered to generate the 4 clusters shown at higher resolution, after filtering to remove cells with high mitochondrial contamination. Functional enrichment terms of each cluster are shown at the right. **H.** Feature plots showing expression of select PCG and lncRNAs marker genes in Kupffer cell subpopulations.

Genes differentially expressed between HFHFD and chow-fed mice at one or more of the above 3 time points were identified for each major liver cell type (Table S4E) and then clustered to give the heatmaps and enriched pathways shown in Fig. 4B-4F. Genes induced across the time course of NAFLD and NASH development were enriched for functions such as fatty acid metabolism and innate immune response in both hepatocytes and endothelial cells (Fig. 4B, Fig. 4C). In HSCs, genes responding to HFHFD were mostly up regulated at both time points and were enriched for cell migration, extracellular matrix, defense response and innate immunity (Fig. 4D), consistent with the activation of NASH-associated fibrosis. Finally, in both Kupffer cells and in NK and T cells, more genes were down regulated than were up regulated, with distinct enriched functions, as shown (Fig. 4E, Fig. 4F).

Several of the lncRNAs identified here as dysregulated with HFHFD feeding have known functions relevant to NAFLD and NASH. These include Pvt1 (lnc12608), which regulates HSC activation to promote liver fibrosis^73, 74^ and was induced 2-fold in endothelial cells from NAFLD liver; and Dnmt3aos (lnc10251), which regulates macrophage polarization^81^ and was induced 16-fold in Kupffer cells (Table S4A). Airn (lnc13654), which promotes hepatocellular carcinoma progression^82^, was induced 5-6 fold in both Kupffer cells and hepatocytes from NASH liver (30 wk). Further, Hnf4aos (lnc1966*), a lncRNA anti-sense to the major hepatocyte transcription factor Hnf4a^83^, was repressed 3-fold in hepatocytes, while lnc-Plet1os (lnc7931), which is anti-sense to a marker gene for epithelial progenitor cells with liver regeneration capacity^84^, was repressed 3-fold in both endothelial cells and hepatocytes (Table S4C). Finally, the oncogenic Snhg8 (lnc2803*)^52^ was down regulated 3-fold in both hepatocytes and Kupffer cells from 34 wk NASH liver (Table S4D).

### LncRNA markers in NASH-associated macrophages

Macrophages play a critical role in NASH pathogenesis; they are strongly linked to disease progression and highly responsive to therapeutic interventions^85, 86^. We re-clustered the Kupffer cells and monocyte/macrophage cells aggregated from both control and HFHFD livers to identify four macrophage subpopulations (Fig. 4G, Table S4G). 96 lncRNA markers were discovered for the individual macrophage subpopulations, of which 27 have human orthologs (Table S4H). The four subpopulations were characterized as Trem2 (low) macrophages, NASH-associated macrophages (NAMs; Trem2-high, a hallmark of mouse and human NASH)^36^, monocyte/monocyte-derived macrophages, and proliferating cells, very similar to those described in^32^ using a non-parenchymal liver cell-enriched population. In NASH liver, Trem2 (low) macrophages decreased from 81% to 30-35% of the overall macrophage population (Table S1H); these cells preferentially express Hotairm1 (lnc5147*), a tumor-associated lncRNA that participates in cell proliferation, migration, and apoptosis^87^, and the Kupffer cell activation marker and scavenger receptor CD163^88^ (Fig. 4H). In contrast, the NAM cell cluster (Trem2-high; Fig. S2) expanded from 8% to 24% of the overall macrophage population in NASH liver and was marked by high expression of Gpnmb (Fig. 4H) and of Dnmt3aos (lnc10251), which regulates macrophage polarization via its effects on the expression of Dnmt3^81^. Monocyte- derived macrophages, which infiltrate and can replace resident Kupffer cells under inflammatory conditions^89^ increased from 5% to 34-40% of the macrophage population (Table S1H), and were marked by Ccr2 and by Mirt1 (lnc15124), an inhibitor of NF-κB signaling that can decrease expression of inflammatory factors^90^. The proliferating macrophage population was marked by Top2a, which is associated with poor prognosis for hepatocellular carcinoma^91^, and by Lockd (lnc5675) (Fig. 4H), which acts as an enhancer of the mitotic cell cycle factor Cdkn1b^92^ and is decreased in offspring liver in response to maternal high fat diet-induced obesity^93^. Other lncRNA markers of macrophage subpopulations are shown in Fig. S2. Analysis of the distinct sets of marker genes for each macrophage subpopulation (Table S4G) revealed unique functions for each cell cluster: Trem2 (low) macrophages were enriched for vasculature development, inflammatory response, response to wounding and cell migration; NAMs were associated with regulation of cell migration/leukocyte migration; monocyte-derived macrophages were enriched for functions related to immune response, inflammatory response, cell death, T cell activation and cytokine production; and proliferating cells were enriched for functions related to DNA replication and cell cycle.

### Zonation dysregulation during NAFLD and NASH pathogenesis

The distinctive liver lobule zone-dependent gene expression seen in healthy liver (Fig. 3) is important for liver function and can be perturbed in disease states^94^. We found significant zonal perturbations in NAFLD and NASH liver compared to chow diet liver for 74 PCGs and 6 lncRNAs in hepatocytes, and for 107 PCGs and 5 lncRNAs in endothelial cells (Fig. 5A, Fig. 5B, Table S4I). The zonally perturbed hepatocyte genes were enriched for pyridoxal phosphate binding and PPAR signaling, while the endothelial cell zonation perturbed genes showed strong enrichment for ribosomal proteins and for NAFLD/respiratory chain (Table S4J). Specific examples include Aldh1a7, which can protect hepatocytes by catabolism of reactive aldehydes formed during oxidative stress and was shifted from pericentral to periportal hepatocytes during NAFLD and NASH development, while the heme biosynthetic enzyme Alas1 shifted from periportal to pericentral hepatocytes (Fig. 5A). In liver endothelial cells, Ssh1 shifted from mid-lobular expression in chow diet livers to pericentral expression in NAFLD and NASH livers (Fig. 5B). This actin remodeling gene is involved in endothelial cell inflammatory signaling and has vascular anti-fibrotic activity^95^.

**Fig. 5.**
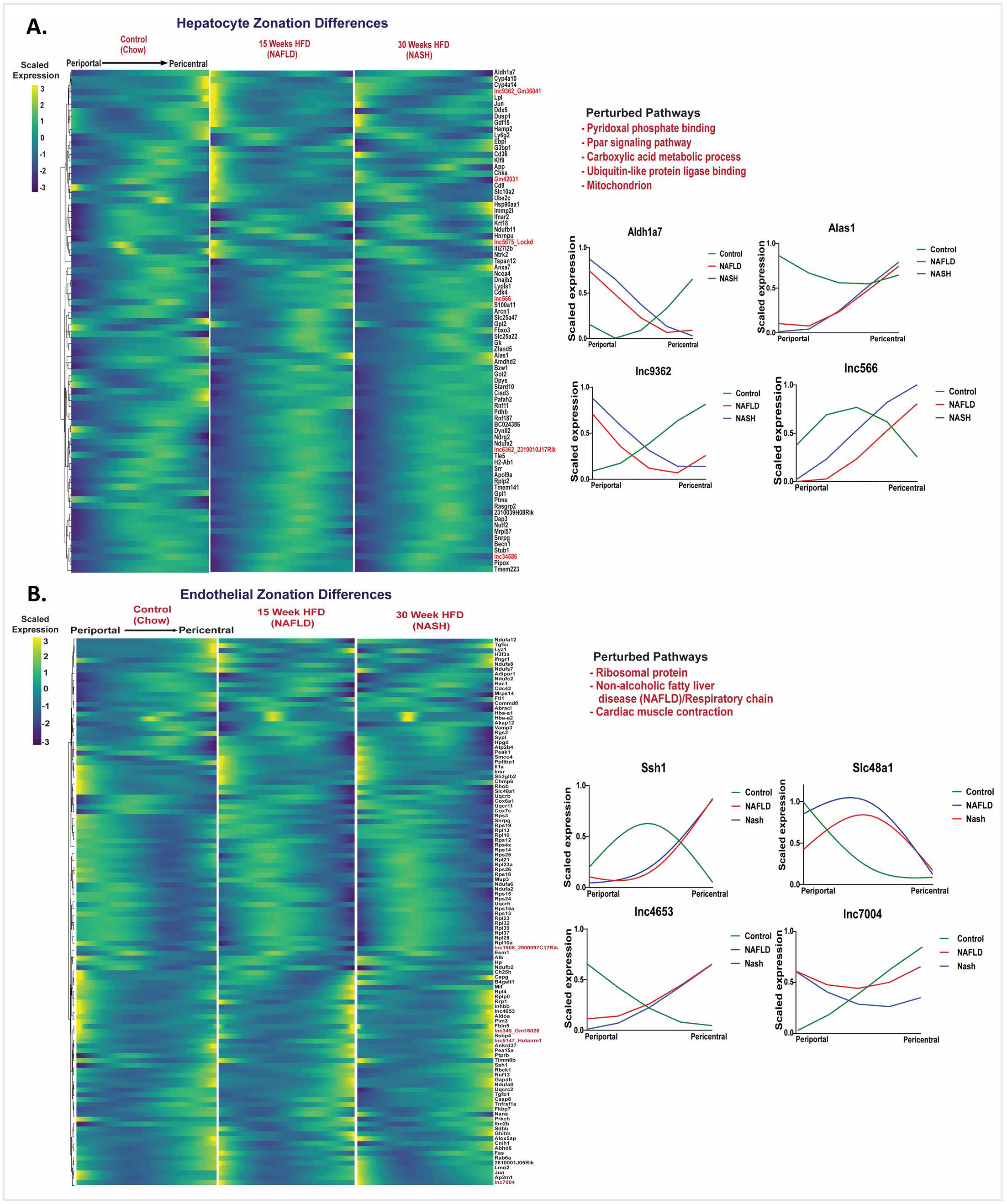
Perturbation of zonation in NAFLD and NASH. Matched heatmaps of genes (PCGs and lncRNAs) that are differentially zonated between control, NAFLD, and NASH livers at FDR <0.001, in hepatocytes (**A**), and in endothelial cells (**B**). Pathways perturbed were identified by DAVID functional enrichment analysis of the differential PCGs (Table S4I, Table S4J). Shown at the right are zonation profiles for select genes for each cell type across three conditions (chow diet, NAFLD, NASH).

### Global discovery of network-essential regulatory lncRNAs in healthy, NAFLD and NASH liver

We implemented gene co-expression network analysis using bigSCale2^96^ to develop gene regulatory networks and associate individual lncRNAs with specific biological functions in healthy liver, in NAFLD liver (15 wk HFHFD) and in NASH liver (30 wk HFHFD) (Fig. 6, Table S5A, Table S5B). We inferred the identities of key network-essential regulatory genes based on 4 network centrality metrics extracted from each network, namely, Betweenness, PageRank centrality, Closeness, and Degree^96^, which serve as proxies for a gene’s influence on the network (see Fig. 6, where labeled nodes are the inferred regulatory genes; Table S5C). A total of 65 such network-essential lncRNAs were identified across the three liver networks. Functional annotation clusters associated with the gene targets of each of these network-essential regulatory lncRNAs (Table S5D) revealed many common enriched functional annotations across the set of lncRNAs. This is consistent with the high gene densities of all three bigSCale2 networks (Table S5E) and the sharing of gene targets between regulatory lncRNAs within a network. Thus, organic acid metabolic process and/or lipid metabolic process described the top two enriched annotation clusters (Benjamini-corrected p-value: E-21 to E-87) for 27 of the 65 regulatory lncRNAs, in either the healthy liver (chow diet) network or the NAFLD network, but not in the NASH network. In contrast, vascular development and closely related functions described the top enriched annotation clusters of 17 other regulatory lncRNAs in the NAFLD and NASH networks but not in the healthy liver network. Finally, regulation of cell migration and/or regulation of angiogenesis described the top two annotation clusters for 12 regulatory lncRNAs in the NAFLD network, but not for the other two networks.

**Fig. 6.**
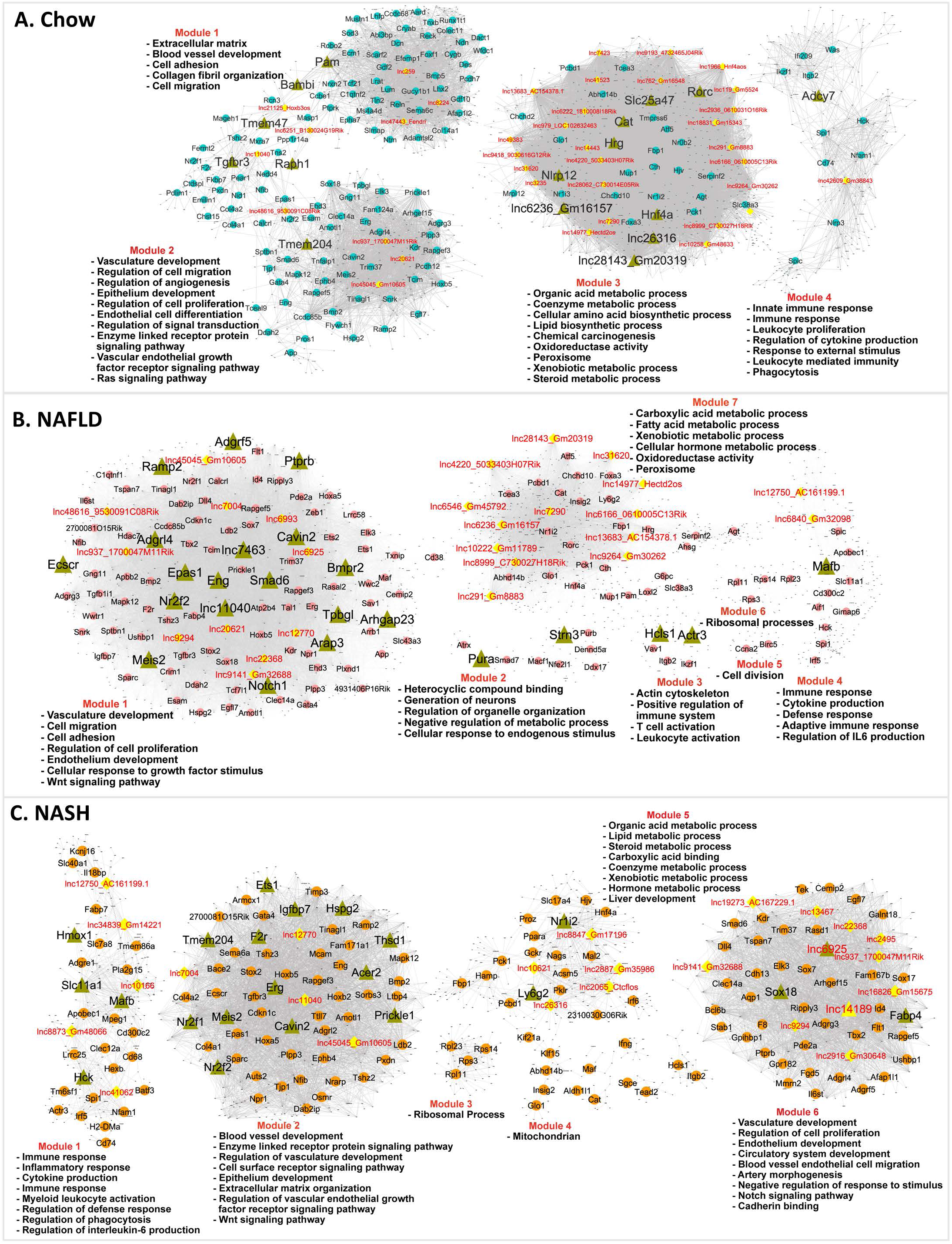
Gene regulatory networks for healthy, NAFLD and NASH liver. BigSCale2 networks based on scRNA-seq data for chow-fed **(A)**, NAFLD **(B)** and NASH **(C)** mouse livers, where PCGs and lncRNAs are nodes and the edges between genes are correlation values based on an adaptive threshold. Nodes displayed here represent network- essential/regulatory genes (circular nodes, with lncRNAs nodes colored yellow), as predicted based on top network metrics. Triangular nodes represent master regulators (predicted to regulate the network-essential regulators) are defined as nodes (genes) with high network centrality metrics calculated for subnetworks extracted from all top 100 ranked PCG nodes plus all top 50 ranked lncRNA nodes. The networks are subdivided into gene modules that are enriched for the biological functions listed. Also see Table S5.

45 genes were identified as network-essential for all three liver networks (Fig. S3; Table S5C, column AH), including 3 lncRNAs, 2 with human orthologs (lnc937*, lnc11040, lnc45045*). Other network-essential regulatory lncRNAs of interest include: Hnf4aos (lnc1966*), which is anti-sense to the major liver transcription factor HNF4A and showed high connectivity to genes involved in various metabolic processes in the healthy liver network (Table S5D); Gm45792 (lnc6546), which is antisense to acyl-CoA synthetase medium chain family member 1 (Acsm1) and is an essential node in the NAFLD liver network; and Ctcflos (lnc2065), which regulates transcription of hepatic Pck1 by modulating glucocorticoid receptor function^97^ and was an essential node in the NASH network, where it makes a second-degree connection with Pck1 via lnc10621 (Fig. S4).

### Master regulators in healthy and NAFLD/NASH liver

The sets of network-essential regulatory genes identified in each network (both lncRNAs and PCGs) were extracted and used to construct subnetworks comprised exclusively of the putative regulatory genes themselves. We ranked the subnetwork genes using a modified scoring approach (see Methods) to identify n=16-23 master regulators for each liver network (Fig. 6, nodes marked with green triangles; Table S5C, columns AP-BU). Seven of the master regulators were lncRNAs (healthy liver network: Gm16157 (lnc6236*), lnc26316, and Gm20319 (lnc28143); NAFLD network: lnc7463, lnc11040; NASH network: lnc6925, lnc14189*). Validating this approach to discovery of bonafide liver network regulatory genes, several of the master regulators are liver transcription factors with well-established regulatory functions, and many of the master regulators have regulatory functions specifically related to liver disease (Table S5F, columns F and L). Specific examples include: Hnf4a, a master regulator of hepatocyte gene expression that protects against NASH development^98^, in the healthy liver network; Nr2f2 (COUP-TFII), whose loss inhibits HSC/myofibroblast activation in liver injury^99^, in the NAFLD liver network; and Meis2, which promotes hepatocellular carcinoma^100^, and Ets1, whose loss decreases diet-induced hepatocyte apoptosis, fibrosis and NASH^101^, in the NASH liver network. We further validated the functionality of the regulatory gene networks by directly comparing the known biological activities of each master regulator to the sets of highly enriched functional annotations of its network target genes (Table S5G), and in many cases found good agreement (Table S5F). For example: Notch1, a master regulator in the NAFLD network, promotes the migration and invasion of hepatocellular carcinoma cells^102^ and its target genes were enriched for regulation of cell migration (Benjamini p=6.46E-17); while the master regulator Cavin2 regulates endothelial nitric-oxide synthase in angiogenesis, and its NAFLD network target genes were strongly enriched for regulation of angiogenesis (p=2.96E-15). Finally, two master regulators from the chow diet liver network that are DNA-binding proteins were identified as upstream regulators of their liver network-predicted target genes by IPA Upstream Regulator analysis: Hnf4a, at 5.80E-19; and Rorc, at 9.71E-08 (Table S5F, column I).

### Cell type-specific responses and lncRNAs dysregulated in CCl_4_-induced liver fibrosis

We examined the utility of the computational framework described above to identify disease-relevant liver lncRNAs in a second disease model. Thus, we analyzed an scRNA-seq dataset^35^ comprised of mesenchymal cells from healthy mouse liver and from livers of mice following chronic (6-wk) exposure to CCl_4_, which induces advanced liver fibrosis. Three distinct cell populations were identified by clustering mesenchymal cells aggregated from the healthy and CCl_4_- treated livers, namely, HSCs, fibroblasts and VSMCs (Fig. 7A, Fig. 7B), consistent with prior findings^35^. Differential expression analysis across these three subpopulations identified 1,550 genes, including 631 lncRNAs, that were dysregulated by CCl_4_ treatment (Fig. 7C, Table S6A), a subset of which were dysregulated in multiple subpopulations (Fig. S6). Genes preferentially or specifically dysregulated by CCl_4_ in the fibroblast subpopulation include a 174-fold induction of Sectm1a, which stabilizes tissue resident macrophages in response to acute inflammation^103^, and a 160-fold suppression of Nkx6-1, an unfavorable prognostic marker for human hepatocellular carcinoma^104^. Genes specifically dysregulated in VSMCs include Stc1, a classic inflammation marker in fibrotic disease^105^, as well as lnc24481*, which was up regulated 16-fold, and lnc43318, which was 5- fold down regulated by CCl_4_ exposure. HSCs showed the most extensive dysregulation, impacting 1,296 genes (743 PCGs, 553 lncRNAs lncRNAs), >90% of which were specifically dysregulated in HSCs (Table S6A). Examples include: Ltbp2 (172-fold increase), a marker of cardiac fibrosis^106^; Gpx3 (16-fold increase), an anti-oxidant enzyme induced by oxidative stress^107^; and Fcna (4-fold decrease), a marker for resting HSCs^108^. Wt1, which moderates fibrogenesis after injury^109^, and the divergently transcribed Wt1os (lnc1601), were both strongly induced (>20-fold) in HSCs. Finally, we identified 172 genes, including 3 lncRNAs, whose expression along the trajectory was significantly different between healthy and chronic CCl_4_-exposed HSCs (Fig. 7D, Table S6B). This finding is consistent with the dynamic changes in zonation seen in HSCs during CCl_4_ -induced fibrotic liver injury^35^. The top enriched term was cell death (Table S6C). In many cases, CCl_4_ exposure substantially reversed the HSC zonation pattern seen in control liver.

**Fig. 7.**
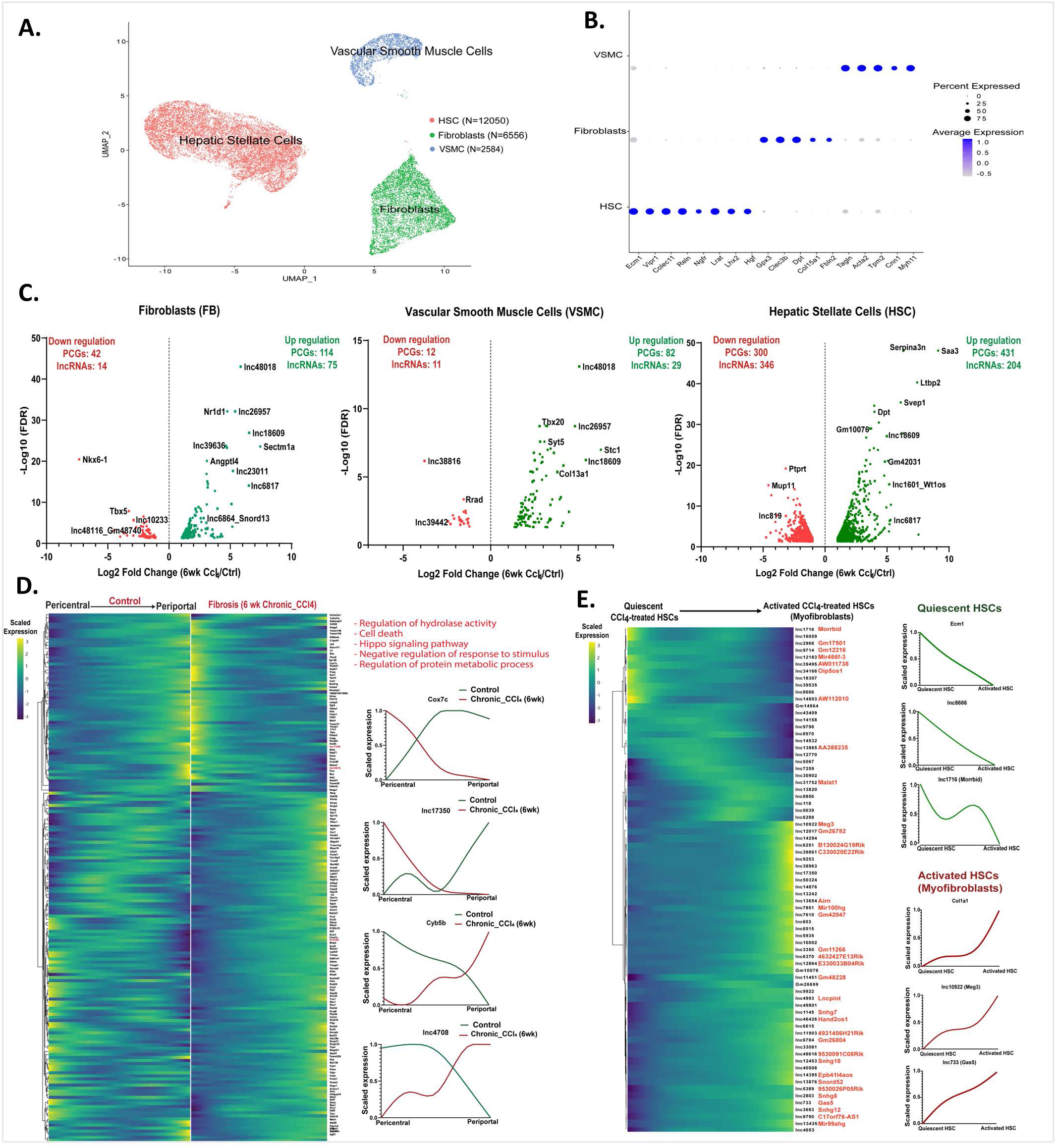
Mesenchymal cell lncRNAs and PCGs perturbed in CCl_4_-induced liver fibrosis. **A.** UMAP showing 21,190 cells comprised of three hepatic mesenchyme subpopulations aggregated from healthy liver (11,710 cells) and fibrotic mouse liver (6-wk CCl_4_ exposure; 9,480 cells), with total cell numbers indicated for each cluster (Table S1H). **B.** Dot plot showing average expression of mesenchymal cell subpopulation marker genes^35^. Dot size is proportional to the percentage of cells expressing each marker gene. **C.** Volcano plots showing differentially expressed genes in each cell cluster at log_2_ |fold-change| >1 and FDR < 0.05, displayed as -log_10_ value on y-axis. **D.** Matched heatmaps of genes expressed in HSCs that are differentially zonated between control and fibrotic mouse liver (at FDR <0.001), with perturbed pathways based on DAVID functional enrichment analysis. *Right*: zonation profiles for select genes whose zonation pattern is perturbed by CCl_4_ exposure. **E.** Heatmap showing relative expression of lncRNAs that are differentially expressed between quiescent pericentral HSCs and myofibroblasts from CCl_4_-exposed mouse liver mesenchymal cells. See Fig. S5B. *Right*: expression patterns for select genes. Cell identities were verified using the uninjured HSC marker gene Ecm1, and the pro-fibrogenic marker gene Col1a1 (right).

### LncRNAs expressed in collagen-producing myofibroblasts

The pericentral HSCs population of CCl_4_-treated liver includes activated HSCs (also known as myofibroblasts), which produce collagen and promote fibrosis progression^35, 108^. We investigated this activated HSC subpopulation to discover lncRNAs associated with pathogenic collagen production. The activation of pericentral HSCs was validated by the expression patterns of pro-fibrogenic marker genes (Col1a1, Col1a2) and by the reduced expression of established marker genes for quiescent pericentral HSCs (Ecm1, Rgs5)^35^ (Fig. 7E, Fig. S5A). Overall, 73 lncRNAs showed differential expression between quiescent and activated pericentral HSCs (i.e., myofibroblasts) from CCl_4_-induced fibrotic liver (Table S6D). For example, the lncRNA Morrbid (lnc1716), a critical regulator of immune cell pro-survival cytokine responses^110^, was more highly expressed in quiescent than in activated HSCs, while Meg3 (lnc10922*), a marker for pulmonary fibroblasts^111^ and an inhibitor of liver fibrosis^25^, and Gas5 (lnc733*), which suppresses HSC activation and counters liver fibrosis^54^, were more highly expressed in activated HSCs. Overall, 1,420 PCGs were differentially expressed between these two pericentral HSC subpopulations (Fig. S5B, Table S6D). PCGs expressed in the quiescent HSCs were strongly enriched for cellular response to chemical stimulus and cell migration (Table S6E), consistent with quiescent HSC migration being linked to cell proliferation^112^, while PCGs in the activated, myofibroblast subpopulation showed strong enrichment for terms such as extracellular matrix, circulatory system development and connective tissue development (Table S6E).

### Network-essential lncRNA regulators in CCl_4_-exposed liver

Analysis of the mesenchymal single cell populations using bigSCale2 yielded functional gene co-expression networks for both healthy (control) and CCl_4_- exposed liver (Fig. 8). Using network centrality metrics, we identified a total of 34 network-essential regulatory lncRNAs: 19 for the healthy mesenchymal cell network and 26 for the CCl_4_-exposed liver fibrosis network; of these, 8 network-essential regulatory lncRNAs were specific to the control liver network, 15 to the CCl_4_ network, and 11 were common to both networks (Table S5C, Fig. S7). One of the network-common lncRNAs, Meg3 (lnc10922*), inhibits HSC activation and accelerates the reversal of CCl_4_-induced fibrosis^113^. Network-essential PCG regulators common to both networks include: Ifnar2, which mediates many anti-viral immune responses and has a gene polymorphism linked to hepatocellular carcinoma^114^; Ets1, which regulates HSC activation in NASH liver and is a profibrogenic marker in CCl_4_ -induced fibrosis^115^, where it is a critical mediator of extracellular matrix remodeling^116^; Tesc, which had the highest PageRank centrality score in the CCl_4_ network and negatively regulates cell proliferation and survival in hepatocellular carcinoma^117^; and Nr1h4 (FXR), which regulates lipid and glucose homeostasis and is involved innate immune responses^118^.

**Fig. 8.**
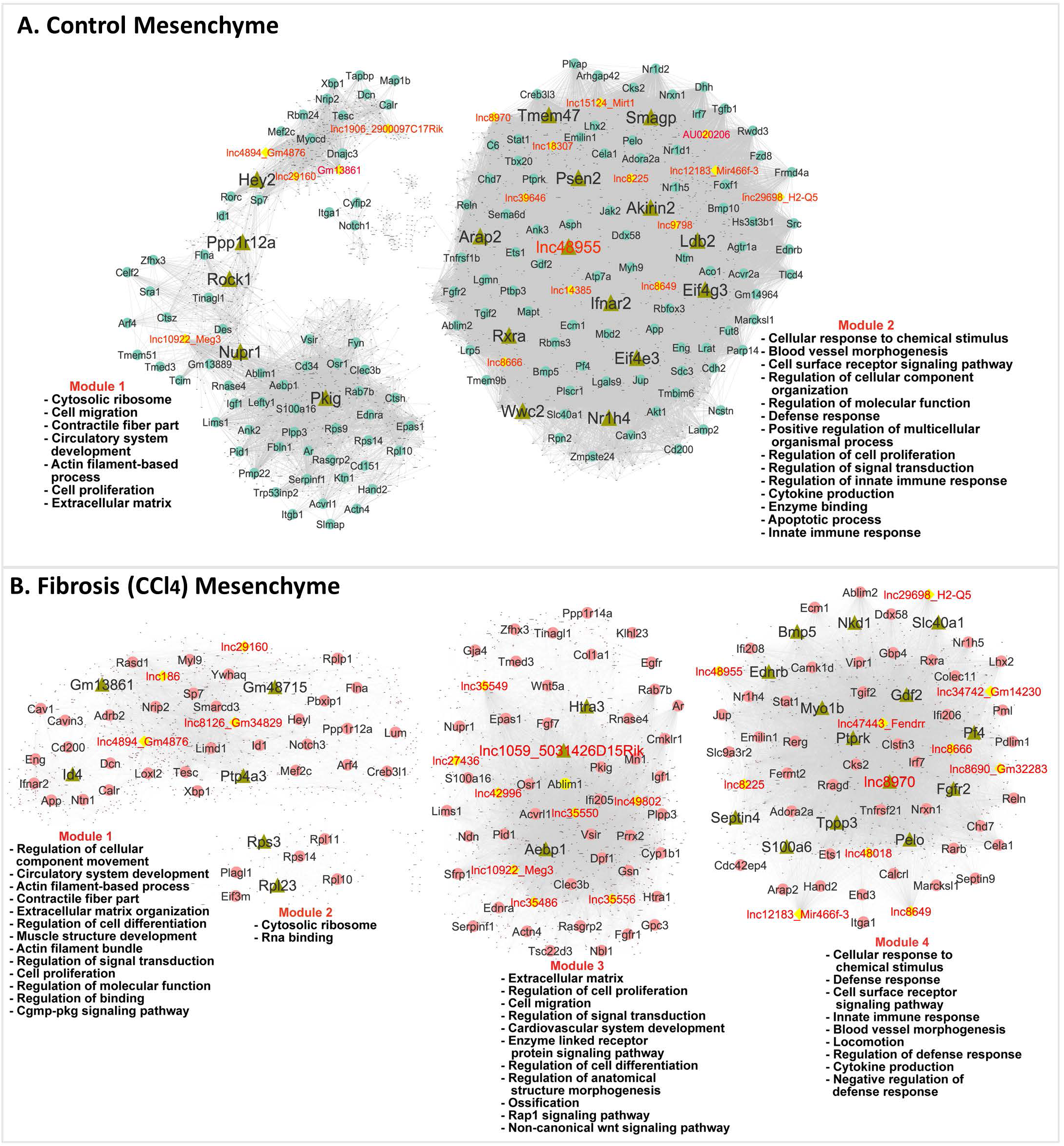
Gene regulatory networks for healthy liver (A) and for CCl_4_-induced fibrotic liver (B). Shown are BigSCale2 networks, with as in Fig. 6, with nodes representing network-essential regulatory genes (circular nodes, with lncRNAs shown as yellow nodes) predicted based on top network metrics. Triangular nodes indicate master regulators. The networks are subdivided into gene modules, which were enriched for the biological functions listed. See Table S5.

Many of the gene targets of the 34 mesenchymal cell network-essential regulatory lncRNAs were enriched for common functional annotations, with notable differences between control and CCl_4_-exposed liver (Table S5D). For example, response to cytokine/innate immune response and related terms were top enriched terms for 14 regulatory lncRNAs in CCl_4_ -exposed liver vs. only 4 regulatory lncRNAs in control liver; and similarly for extracellular matrix (9 regulatory lncRNAs in CCl_4_ liver vs. 1 in control liver), contractile fiber/actin/muscle system process (6 lncRNAs in CCl_4_ liver vs. 3 in control liver) and locomotion (7 lncRNAs in CCl_4_ liver vs. 11 in control liver). These findings are consistent with the central roles of each of these biological processes in liver fibrosis. In contrast, top enriched terms more frequently found for gene targets of regulatory lncRNAs from control liver included response to organic substance (9 in regulatory lncRNAs in control liver vs. 0 lncRNAs in CCl_4_ liver) and blood vessel/cardiovascular system development (13 lncRNAs in control liver vs. 8 lncRNAs in CCl_4_ liver) (Table S5D).

### Master regulators in the mesenchymal cell networks

Analysis of the sets of mesenchymal cell network regulatory genes as described above for healthy and NAFLD/NASH identified n=18-23 master regulators for each mesenchymal cell network along with enriched functional annotations of their target genes (Fig. 8, nodes shown as green triangles; Table S5C, Table S5F). Four lncRNAs were identified as master regulators of the CCl_4_ mesenchymal network: Gm13861 (target genes enriched in cytoskeletal protein binding, at Benjamini p=7.12E-10); Gm48715 (muscle system process, p=9.39E-09), lnc1059* (extracellular matrix, p=7.64E-11); and lnc8970 (blood vessel morphogenesis, p=1.66E-11). One master regulatory lncRNA was identified in the control mesenchymal network (lnc48955, targets enriched in locomotion, at p= 4.62E-22) (Fig. 8, Table S5C, Table S5F, Table S5G). Validating our approach to discovery of liver network-essential regulatory genes, many of the master regulators from the CCl_4_-exposed mesenchymal cell network have regulatory roles in liver fibrosis, a hallmark of CCl_4_-exposed liver, and the network target genes of the master regulators often showed highly enriched functional annotations that match the known biological properties of their master regulators (Table S5F).

### Functional clustering of regulatory lncRNAs

PCGs that were targets of the network-essential regulatory lncRNAs from each of the above 5 liver networks (Fig. 6, Fig. 8) were input to Metascape^119^ to facilitate comparisons across the target gene lists of each lncRNA, and thereby cluster the regulatory lncRNAs based on commonality of function. Regulatory lncRNAs from chow fed liver comprised three major functional clusters (Fig. S8), two of which were variously enriched for diverse metabolic processes, while a third lncRNA cluster was functionally enriched for angiogenesis and vasculature development, extracellular matrix, and cell-cell adhesion. Similarly, we identified three clusters of regulatory lncRNAs from the NAFLD network (Fig. S9), and four clusters with distinct enrichment patterns for the regulatory lncRNAs from NASH liver (Fig. S10). Overall, the NAFLD and NASH network-derived regulatory lncRNAs showed more extensive enrichment for vasculature development and angiogenic processes than those from control liver (Fig. S11), whereas the NAFLD and control network regulatory lncRNAs both showed enrichment for diverse metabolic processes.

Similarly, we identified two distinct regulatory lncRNA clusters from the healthy liver mesenchymal cell network (Fig. S12), with one cluster of 5 lncRNAs enriched for terms related to muscle contraction, notch signaling, and ion homeostasis, and a separate cluster comprised of 14 lncRNAs enriched for diverse biological processes, including cell morphogenesis, regulation of defense response, vascular development, and interferon responses. Finally, regulatory lncRNAs from the CCl_4_-exposed mesenchymal cell network yielded three clusters with strongest enrichments and highest specificities for muscle contraction and notch signaling, extracellular matrix organization, and response to interferon-beta, respectively (Fig. S13). Pathways that were either common or specific to healthy versus CCl_4_-exposed mesenchymal cell lncRNA gene targets are shown in Fig. S14.

### Triplex potential of liver disease-associated regulatory lncRNAs

Finally, we evaluated the potential of each of the above network-essential regulatory lncRNAs to form a triple helix with the promoter sequence of its putative protein coding target genes in the gene co-expression network, as determined by Triplex Domain Finder^120^. Specifically, for each IncRNA, we computed the enrichment of sequence-specific lncRNA-protein coding gene promoter triplex interactions in the IncRNA’s network-predicted target gene set, compared to a background set of protein coding gene promoters. First, we used Triplex Domain Finder to define a set of DNA-binding domains within each lncRNA of interest, i.e., lncRNA sequences with high triplex forming oligonucleotide activity. We then determined, for each such DNA-binding domain, whether the set of target gene promoters was significantly enriched for containing at least one Triplex Target Site (TTS) as compared to that for the set of non-target gene promoters. For the chow-fed liver network, we determined that 7 of 39 lncRNAs identified as key regulatory genes based on network metrics (Table S5C) showed significant enrichment for lncRNA-promoter DNA triplexes at FDR <0.05 (Fisher’s exact test) (Table S7A). Four of these 7 lncRNAs formed triplexes with genes enriched for metabolic processes such as oxidoreductase, peroxisomes and lipid metabolism (Fig. S15A; 5033403H07Rik (lnc4220), lnc7423, 0610031O16Rik (lnc2936), Prox1os (lnc979*)) and 3 lncRNAs formed triplexes with genes enriched for extracellular matrix (Table S7A; lnc259, Fendrr (lnc47443*), lnc8224). Furthermore, 9 out of 28 regulatory lncRNAs in the NAFLD network were significantly enriched for triplex formation, and 5 out of 29 regulatory lncRNAs in the NASH network were significantly enriched for triplex formation with their gene targets. Enriched gene functions associated with these regulatory lncRNA gene targets include vasculature development and angiogenesis (Fig. S15B, Fig. S15C).

In healthy liver mesenchymal cells, 8 out of 19 network-essential regulatory lncRNAs were significantly enriched for triplex formation with their target gene promotors; the associated enriched functions included response to chemical stimulus, vasculature development, cell migration, immune response, and extracellular matrix (Fig. S16A). In the CCl_4_-induced liver fibrosis network, 11 out of 26 regulatory lncRNAs were enriched for triplex formation with their target gene promoters for pathways linked to extracellular matrix, response to cytokine and interferons (beta and gamma), blood morphogenesis, and defense response (Fig. S16B). Six network- essential regulatory lncRNAs were common to both the control and the CCl_4_-exposed liver mesenchyme network, but new regulatory interactions emerged in the CCl_4_ network. For instance, Meg3 (lnc10922*) formed an isolated subnetwork in control mesenchymal cells (Fig. S16A) but had extensive shared binding targets with lnc35550*, lnc35556* and 5031426D15Rik (lnc1059*) in the liver fibrosis network (Fig. S16B). We identified specific PCG targets for each lncRNA, as well as many shared targets between regulatory lncRNAs, which is indicative of their complex regulatory crosstalk (Tables S7C-S7G).

## Discussion

The overall goal of this study was to elucidate on a global scale the roles of liver-expressed lncRNAs in biological pathways related to liver disease development. Single-cell RNA-seq was employed to characterize the long non-coding transcriptional landscape of mouse liver using a reference catalog of 48,261 liver-expressed lncRNAs, a majority of them novel, which we discovered by transcriptome reconstruction from > 2,000 bulk public mouse liver RNA-seq datasets, a major update to our earlier reference set of 15,558 mouse liver lncRNAs^13^. Prior studies identified several hundred mouse liver-expressed lncRNAs that show hormone-regulated, sex-biased expression^13, 121^ or respond to xenobiotic exposures^16, 17, 19^. However, a global analysis of lncRNAs dysregulated in liver diseased states, most notably, lncRNAs expressed in specific cell types and zonal subpopulations across the liver lobule, was lacking. Here, we characterized liver cell type-specific expression patterns for healthy mouse liver and for two disease models for a total of 76,011 genes, including 48,261 liver-expressed lncRNAs and 4,700 other lncRNAs from RefSeq and Ensemble databases. Importantly, we showed that single cell RNA-seq technology is sufficiently sensitive to detect and characterize more than 30,000 liver lncRNAs, including 25,000 novel liver-expressed lncRNA genes, 110 of which we identified as cell type-specific marker genes for the 13 major cell types identified in healthy adult mouse liver. Using public scRNA-seq datasets^33, 35^ we uncovered marked liver cell type-dependent perturbations in the expression of 677 lncRNAs in NAFLD compared to healthy mouse liver, and in the transition from NAFLD to NASH liver. Furthermore, we identified a largely non-overlapping set of 631 lncRNAs dysregulated in liver mesenchymal cells in liver fibrosis induced by chronic CCl_4_ exposure. Given that thousands of liver-expressed lncRNAs are nuclear and tightly bound to chromatin^19^, we can expect an even greater sensitivity for lncRNA detection when using single nucleus RNA-seq in place of scRNA- seq^15^. Single nucleus RNA-seq may also provide a more representative picture of the relative abundance of each cell type in mouse liver, something that is lacking in the integrated UMAP of healthy (normal chow diet) mouse liver single cell populations used in our analysis (Fig. 2), which were aggregated and harmonized across four datasets enriched for mouse non-parenchymal cells.

Single cell analysis has enabled the functional characterization of the periportal to pericentral zonation gradients of PCG expression across the liver lobule, as seen in hepatocytes^30^, endothelial cells^40^ and HSCs^35^. Here, we used spatial inference methods to elucidate zonation patterns for large numbers of lncRNAs, and PCGs, in each of five major liver cell types (hepatocytes, endothelial cells, HSCs, fibroblasts and VSMCs). The expression profiles reported here for VSMCs and fibroblasts are novel; prior reports were limited to extrahepatic tissues, including brain^77, 78^, whose zonated markers we used to annotate the VSMC zonation clusters identified in liver. Liver fibroblasts were subclustered to give two novel subtypes of unknown function, one of which was uniquely marked by Colec11, which plays an important role in innate immunity^122^, and the other by Ly6c1, an antigen expressed by monocytes/macrophages that ingest stressed erythrocytes and deliver iron to hepatocytes^123^. Further study, including experimental validation by single-molecule fluorescence in situ hybridization, will be needed to verify these zonated lncRNA expression patterns.

Single cell analysis can dissect the milieu of hepatic immune cells^32^, which are critical for hepatic immune surveillance and immune tolerance^124^ and protect against pathogens and dietary antigens^125^. Kupffer cells (liver resident macrophages) are essential for tissue repair and clearance of toxins^126^ and can differentiate into the more highly pathogenic NASH-associated macrophages (NAMs)^32^. We identified a total of 96 lncRNA markers for four macrophage subpopulations from NAFLD and NASH liver, some of which may be useful as clinical indicators for disease diagnosis or progression and perhaps serve as functional therapeutic targets^23^. These four cell subpopulations are consistent with prior findings^32^ and include Trem2-low macrophages, which are involved in innate immunity, NAMs (Trem2-high cells), which are an indicator of disease progression and provide opportunities for therapeutic intervention^127^, and monocyte-derived macrophages, which are responsible for chemokine-mediated signaling and leukocyte migration and may replace the depleted Kupffer cells^126^. Finally, proliferating Kupffer cells were characterized by high expression of cell division and cell proliferation genes (e.g., Top2a, Stmn1; Fig. S17) and reflect the increase in macrophage proliferation in response to damage induced by NAFLD and NASH.

We explored lncRNA responses in chronic CCl_4_-exposed liver, a widely used model for chemical-induced liver fibrosis and identified 631 lncRNAs dysregulated in or more hepatic mesenchymal cell populations. Validating this model, we observed strong up regulation of many HSC-expressed lncRNAs in association with the up regulation of PCGs involved in extracellular matrix and collagen processes, a key feature of HSC activation and liver fibrosis^128^. Further, spatial inference analysis revealed apparent zonation differences between control and fibrotic HSCs. Importantly, we identified 73 lncRNA markers, many of them novel, for the transition from the quiescent HSC state to the activated state of collagen-producing HSCs (myofibroblasts) following CCl_4_-induced centrilobular injury. Future studies may investigate these lncRNAs as potential fibrosis biomarkers and therapeutic targets and may expand the analysis reported here to include effects of CCl_4_-induced hepatotoxicity on the full repertoire of liver cell types.

Gene regulatory network inference is a promising approach to discover regulatory mechanisms based on transcription factor-target gene interactions in both healthy tissue and in disease phenotypes. We utilized the rich single cell datasets analyzed here to construct gene regulatory networks, with three specific goals: 1) to deduce lncRNA function based on membership in a functional gene module; 2) to identify network-essential regulatory lncRNAs, i.e., lncRNAs that are expected to be critical for specific linked biological pathways; and 3) to discover new regulators that emerge in the rewiring of networks in diseased states. We analyzed the bigSCale2- derived networks obtained for both healthy and diseased liver (NAFLD, NASH, and fibrotic liver) using four distinct network centrality metrics to discover network-essential regulatory genes, including many lncRNAs. The four network metrics used capture different types of network regulators. Thus, Betweenness is crucial for information flow between network modules, Closeness identifies genes that can rapidly spread information across the network, PageRank centrality marks genes that are highly influential on the network, and Degree, a measure of a node’s centrality, identifies key network hubs^96^. Master regulators, both lncRNAs and PCGs, were also identified for each network based on their central roles in subnetworks comprised exclusively of the network-essential regulatory lncRNAs and PCG genes themselves. Many of the regulatory lncRNA gene targets from the diseased liver networks were enriched for common sets of functional annotations that are characteristic of the diseased states. For instance, clusters of lncRNAs in the NAFLD network were associated with cell migration and regulation of angiogenesis, while the sets of NAFLD and the NASH network regulatory lncRNA target genes were both associated with vascular development, and in CCl_4_-exposed mesenchymal cells, with cytokine and innate immune response and extracellular matrix. Validating this approach to the discovery of network regulatory genes, many of the master regulator PCGs that we identified have known regulatory functions specifically related to liver disease, and many of their network target genes were highly enriched for functional annotations that match the established biological functions of their master regulators.

LncRNAs can act as scaffolds that promote interactions between proteins, RNA, and DNA^129^. There is increasing evidence that some lncRNAs interact with genomic DNA in a sequence-specific manner via triple helix (triplex) formation, which may enable lncRNAs to assemble protein complexes at specific genomic regions and regulate gene expression^120, 130^. We used the sequence-based algorithm Triplex Domain Finder to predict RNA–DNA triple helix binding between regulatory lncRNAs identified in our bigSCale2 networks and the DNA promoter sequences of the genes that make direct connections with those lncRNAs in the gene regulatory network. We found significant enrichments of triplex interactions in the set of target genes for 20-32% of the network- essential regulatory lncRNAs identified in the NAFLD and NASH networks and for 42% of those from the CCl_4_- induced liver fibrosis network (Table S7A). Little overlap was found between the NAFLD and NASH network- essential regulatory lncRNAs that formed significant triplexes with PCG promoters and those from the CCl_4_- induced fibrosis network. This is not unexpected as the fibrosis network input was limited to single cell expression data from hepatic mesenchymal cell populations. Finally, within networks, we observed many shared gene targets between regulatory lncRNAs, which indicates substantial regulatory crosstalk (Fig. S15-S16).

In conclusion, we present a comprehensive characterization of thousands of novel mouse liver lncRNAs including gene and isoform structures, cell-type expression patterns, inferred spatial location across the liver lobule, and dysregulation in liver disease. Many novel lncRNAs were identified as markers for pathogenic cell types in NASH and in liver fibrosis. A computational framework implemented to discover network-essential regulatory lncRNAs and their predicted gene targets was validated by multiple examples drawn from network-essential regulatory PCGs of known function. Finally, for a subset of the putative regulatory lncRNAs that we identified, promoter sequences upstream of the network-defined lncRNA target genes were shown to be significantly enriched for IncRNA triplex formation, a finding that gives independent mechanistic support for the lncRNA−target genes linkages predicted by our gene regulatory networks.

### Limitations of the study

As this study is based on scRNA-seq data, there are technical limitations related to the sparse nature of such data sets, in particular when it comes to characterization of lncRNAs whose overall expression in the cell is often much lower than that of PCGs; greater sensitivity for lncRNA detection would likely have been achieved by using single nucleus RNA-seq. Further, our analyses were limited to the specific set of 48,261 liver-expressed lncRNAs that we identified based on our analysis of more than 2,000 bulk mouse liver RNA samples, plus 4,700 other lncRNA genes that we collected from established databases. The absence of expression data for other, presently unknown lncRNAs that may be important for liver function and pathophysiology is another limitation. Several of our conclusions are based on inferences drawn from analyses that are, at their core, correlative in nature. These include our analysis of pseudo-time trajectories to infer zonation patterns for PCGs and lncRNAs across the liver lobule, as well as changes in zonation in liver disease. While such changes in zonation have, in some cases, been verified by others experimentally for certain PCGs, the lack of experimental verification of the lncRNA zonation patterns reported here is a limitation. Furthermore, while we were able to validate our use of single cell-based gene regulatory networks to identify network- essential genes, as evidenced by multiple examples of network-essential regulatory PCGs whose known biological functions align with the enriched functions of their gene targets in the regulatory networks, the inferred nature of the gene regulatory networks is nevertheless a limitation. Experimental validation of the network-essential lncRNAs described here using lncRNA knockout or knockdown mouse models will ultimately be required to address this limitation.

## Author contributions

KK and DJW jointly conceptualized the study and its major aims. KK carried out all of the computational analysis, including methodology development, script writing and figure preparation. Downstream data analysis and supplementary table preparation were carried out by KK with the assistance of DJW. KK prepared a draft manuscript and DJW revised and edited the manuscript and finalized the text and supplementary tables. DJW was responsible for project funding acquisition, project supervision and administration.

## Acknowledgements

Supported in part by NIH grant ES024421 (to DJW).

## Methods

### Transcriptome reconstruction

We downloaded from GEO, ENA and Array express a total of 2,089 bulk mouse liver RNA-seq samples extracted from 68 different studies (Table S1A), all sequenced by paired-end, stranded Illumina RNA sequencing. Sequence reads were mapped to the mouse mm9 reference genome using Tophat2, and Cufflinks was used to assemble mapped reads into transcripts for each individual RNA-seq sample. We then used two algorithms for transcriptome assembly: TACO^43^ and Cuffmerge^44^, each of which produced a different assembly comprised of both coding and noncoding transcripts. The TACO-derived transcriptome was evaluated for read-through transcripts^131, 132^, a product of transcriptional read-through of two adjacent genes on the same strand, which in many cases represent spurious transcriptional noise. 23,346 out of 1,491,436 TACO-assembled transcripts were identified as potentially read-through based on their overlap with multiple PCGs or lncRNAs. For each of these putative read-through transcripts, we calculated isoform_sum_abs_frac, that is, the absolute fraction of all transcript isoforms of a given gene that is represented by the sum total of all the read-through isoforms for that gene. Transcripts whose isoform_sum_abs_frac was > 0.1 were retained, and the associated genes concatenated to give a new, longer gene structure. For transcripts whose isoform_sum_abs_frac was ≤ 0.1, we discarded the entire set of read-through transcripts for that gene, as they likely represent transcript assembly artifacts. Overall, 19,219 (82%) of the 23,346 read-through transcripts initially identified were discarded. In parallel, we assembled the transcriptome of each of the 2,089 RNA-seq samples with Cufflinks (v2.1.1)^44^ and discovered 547,300 transcripts that were used for downstream filtering and lncRNA discovery.

### lncRNA discovery

The filtered transcriptome obtained from each assembly method (TACO and Cufflinks) was further processed using two different lncRNA discovery pipelines, as we described previously for rat liver lncRNAs^18^. <u>Method 1</u>: LncRNA transcripts were identified based on transcript length > 200 nt, low or no coding potential, and absence of overlap with known PCGs^14^. <u>Method 2</u>: the lncRNA discovery tool Slncky^45^ was used to remove transcripts that overlap PCGs, and to assess the coding potential for small peptides and novel proteins.

A filter based on synteny was used to remove transcripts that align to syntenic coding transcripts in other, related species. A non-synonymous to synonymous (dN/dS) ratio was then calculated and used to evaluate the coding potential of each transcript. The TACO assembly yielded 109,937 sequences that passed our filters to qualify as lncRNAs but were found to be mono-exonic fragments that overlap a PCG intronic region. These were filtered out as likely derived from spurious transcription, and consequently, no mono-exonic intragenic lncRNAs were included in the final TACO assembled lncRNA gene list. Bedtools (v2.17.0) was used to perform overlap analysis between the set of 44,579 lncRNA gene structures obtained by TACO assembly, the 25,869 lncRNA gene structures generated by Cuffmerge, and the original set of 15,558 liver-expressed lncRNAs that we discovered previously using a much smaller set of input RNA-seq samples^13^. These lncRNA genes and their isoforms were merged using the Bedtools overlap command. The gene structures and isoforms in the original set of 15,558 liver-expressed lncRNAs were updated based on the TACO and Cuffmerge assemblies while retaining the original lncRNA gene numbers^13^. In many cases, the new lncRNA gene structure represents a merging of two or more adjacent lncRNAs from the original 15,558 lncRNA gene list; those lncRNAs were renamed by concatenating the old lncRNA gene numbers while using the lowest number to represent the new lncRNA structure. For example, the new lnc_inter_chr8_7423 (chr8:116,533,047-116,655,614) was obtained by merging the prior set of tandem minus-strand lncRNAs, numbered lnc7423 and lncs 7425-7430, and consequently, there are no lncRNAs numbered 7425 through 7430 in the final set of 48,261 liver-expressed lncRNAs. Lnc_inter_chr8_7424 (chr8:116,533,910-116,537,738) is a plus-strand lncRNA that is intragenic and anti-sense to the new lnc_inter_chr8_7423 and was therefore not included in the merger that formed the new lnc_inter_chr8_7423 (Table S1B). Newly discovered lncRNAs that did not overlap the prior set of 15,558 lncRNAs were assigned new lncRNA gene numbers, beginning with lncRNA number 15,559 on Chr1.

### Final set of 48,261 liver-expressed lncRNAs (mm10)

The final set of 48,361 lncRNAs and their total of 150,280 isoforms, defined for mouse genome mm9 (Table S1B, Table S1D), was reduced to 48,261 lncRNAs after lift-over to mouse genome mm10 (Table S1C), and was comprised of the following (based on mm10 mapping) : 1,176 *intragenic lncRNAs*, which overlap one or more PCG intronic regions on the same strand, and whose exons do not overlap any PCG exonic regions on the same strand; 4,127 *antisense lncRNAs*, which overlap a PCG on the opposite strand; 42,892 *intergenic lncRNAs*, which do not overlap any PCG on either strand; and 66 *intra- antisense lncRNAs*, which have attributes of both antisense and intragenic lncRNAs. 42,605 of the 48,261 lncRNAs are mono-exonic genes and 5,656 are spliced, multi-exonic genes. The designation ’mono-exonic’ is used here to describe a lncRNA whose union of all exons across all isoforms (’exon collapsed’ sequence) covers the entire gene body without leaving any intronic gaps. However, 7,754 of the 42,605 mono-exonic lncRNAs have multiple isoforms, which in many cases include intronic sequences (Table S1D), and thus could be considered multi-exonic.

The *number* of lncRNAs referred to in this study, e.g., the number of IncRNAs expressed in a given liver single cell cluster, or the number of lncRNAs that are differentially expressed under a given biological condition, is based on a total lncRNA count of 52,961, corresponding to the subset of the 48,261 liver-expressed lncRNAs that meet the specified condition, plus the subset of 4,700 other long-noncoding genes that met the same condition: 4,697 Ensembl noncoding lncRNAs assigned the gene biotype ‘lincRNA’ (N=3,176) or ‘anti-sense’ (N=1,521), plus 3 other lncRNAs not included in the above lists (lnc-LFAR1, LeXis, Lnclgr) (genes listed in Table S2A).

### Ortholog conservation

We employed Slncky^45^ to assess primary sequence conservation and syntenic conservation to discover IncRNA orthologs in the human and rat genomes for the full set of 52,961 mouse liver lncRNAs described above. Two metrics were used to identify orthologs, as implemented by Slncky: transcript- genome identity (TGI; percent of identical, aligning sequences between a mouse lncRNA and a non-transcribed locus in a syntenic region in the rat or human genome); and transcript-transcript identity (TTI; percentage of identical, aligning sequences present in the exonic regions of lncRNAs in both species)^45^. A threshold of 30% conservation based on either the TTI metric or the TGI metric was used to define an orthologous human or rat sequence. To identify human orthologs, we input to Slncky a set of 18,151 human lncRNAs from GENCODE (version 23)^133^, 96,308 human lncRNAs from NONCODE (v5)^134^, and 662 human lncRNAs involved in cancer (onco-lncs) that we curated from PubMed. To identify rat orthologs, we input our published set of 5,795 rat liver lncRNAs^18^. As this set of rat lncRNAs is far from complete, there are likely to be many more than the 1,805 rat lncRNA orthologs identified here (Table S1E, Table S1F). We found human orthologs for 9,543 of the 48,261 mouse liver-expressed lncRNAs (Table S1E) and for 1,166 of the 4,700 other non-coding lncRNAs described above (Table S1F).

### GTF file for single cell and single nucleus RNA-seq analysis

We created a custom reference GTF file for the mouse mm10 genome build for use with the CellRanger (v3.1.0) mkref command^135^. This GTF file is comprised of gene coordinates for 76,011 genes (Table S2A) plus 91 ERCC spike-in controls: RefSeq PCGs (N=20,973; 19,801 genes with NM accession numbers, 1,159 genes with both NM and NR accession numbers, and 13 mitochondrial genes), the above-described set of mouse liver lncRNAs with mm10 genome coordinates (N=48,261), RefSeq non-coding genes defined by NR accession numbers that do not overlap the set of 48,261 lncRNAs (N=2,077), Ensembl noncoding lncRNAs that do not overlap RefSeq NR genes or the set of 48,261 lncRNAs (N=4,697; 3,176 genes designated ‘lincRNA’ and 1,521 genes designated ‘anti-sense’ in Table S2A), and three other lncRNAs (lnc- LFAR1, LeXis, Lnclgr) absent from the above lists. The full set of mouse liver lncRNAs identified in our analysis, originally discovered by mapping bulk RNA-seq fastq files to the mm9 reference genome, was converted to mm10 genomic coordinates using the UCSC genome browser LiftOver tool^136^. The GTF file used for scRNA-seq analysis (gtf76k_mm10_Liver_scRNAseq.gtf, available in Supplementary materials) contains exon region gene coordinates for all 76,011 genes and was used by CellRanger to count sequence reads mapping to exons.

### Datasets and data processing for single cell analysis

Single-cell RNA-seq data for mouse liver was downloaded from GEO (https://www.ncbi.nlm.nih.gov/geo/). Raw sequencing data for hepatocytes (GSE109774)^137^ and for the liver non-parenchymal cell clusters shown in Fig. 2A (GSE137720, E-MTAB-8077, and GSE129516)^32, 35, 42^ was obtained from healthy (untreated, normal chow diet) mouse livers (Table S1G). We also analyzed 10x Genomics scRNA-seq datasets from GSE166504^33^ to study the effects of high fat diet-induced NAFLD and NASH, and from GSE137720^35^ to study CCl_4_-induced liver fibrosis (Table S1G). Raw fastq files were processed and aligned to the mm10 mouse reference genome and processed using the single cell analysis 76,011 gene custom GTF file described above. Feature-barcode matrices were generated using CellRanger software (v3.1.0)^135^. Multiple scRNA-seq datasets were combined using the CellRanger aggr command and count data were then processed using Seurat v3^138^.

### Integration and clustering

Using Seurat v3, we normalized the integrated CellRanger count matrix data by dividing the UMI count per gene by the total UMI count in the corresponding cell followed by log-transformation. Cells with fewer than 200 genes detected, fewer than 400 UMI per cell, or cells with >5% gene reads mapping to the 13 mitochondrial genome genes were excluded from all downstream analyses. Highly variable genes were identified using the FindVariableGenes function of Seurat with default parameters. Variable genes were projected onto a low-dimensional space using principal component (PC) analysis, with the number of PCs set to 10 based on an inspection of elbow plots of the variance explained for each dataset. We used Harmony (v1.0)^139^ to remove from the embedding the influence of dataset-of-origin factors (e.g., the effects of varying batches or technologies) and to integrate shared cell types across different datasets. We input to Harmony normalized gene matrix files saved as a Seurat object along with a pre-calculated PC analysis embedding based on 10 PCs using default parameters. Graph-based clustering was performed to cluster the cells based on their gene expression profiles using the *FindClusters* function in Seurat. The clusters obtained were visualized using the UMAP function of Seurat v3 (resolution: 0.15 and PC=10), and cell identities were assigned to each cluster based on the expression of established marker genes (Fig. S1). scDblFinder^140^, a doublet detection algorithm, was applied (with default parameters) to identify and remove doublets based on artificial doublet generation, parameter optimization and thresholding.

### Detection of marker genes

Cell-type specific marker genes that drive the separation between single cell clusters were identified by pairwise differential gene expression analysis ^141^, comparing each cell cluster against all other cell clusters using the Global Distinguishing function of Loupe Browser (10x Genomics, Loupe Browser 5.0 (https://www.10xgenomics.com/). Genes expressed in >5% of the cells in a cluster were identified as putative marker genes for that cluster if they showed differential expression at log2 fold-change >2 and FDR (Benjamini-Hochberg) < 0.05 when compared to the set of all cells comprising all other clusters. We then removed from the list of putative marker genes all genes that were identified as putative marker genes for two or more clusters, resulting in the final set of cluster-specific marker genes (Table S2D). Marker genes distinguishing the 4 macrophage clusters shown in Fig. 4G were identified using a relaxed fold-change filter of log2 |fold-change| >1 because of the close relationship between the 4 cell populations.

### Zonation analysis

Pseudo-temporal trajectories were inferred from the hepatocyte population obtained from chow-fed mouse liver^33^ and for each of four major non-parenchymal cell clusters (endothelial cells, HSCs, fibroblasts, VMSC; Fig. 2A) using the R package Monocle2 (v2.6.419)^142^ with default parameters. Cells were ordered based on the inference pseudotime trajectory in Monocle2. PCGs and lncRNAs showing significant changes in expression along the pseudotime trajectory were identified using generalized linear models (differentialGeneTest function in Monocle 2) with a threshold of q-value < 10^-3^ for significance. Hierarchical clustering of the genes that were co-zonated across pseudotime was implemented using the *pheatmap* function of Monocle2. The zonation clusters were annotated as periportal versus pericentral for hepatocytes^41^, endothelial cells^42^ and HSCs^35^ using established marker genes, and the resulting pseudotime values were scaled between 0 and 1. Cell trajectories were also determined for HSCs from healthy and from CCl_4_-treated livers. Hierarchical clustering in Monocle2 divided zonation heatmap s into clusters, which were assigned labels based on established zonation marker genes in each major cluster. SCORPIUS^70^ was used to further validate the endothelial cell zonation patterns (Fig. 3B). We included traditional endothelial cell phenotypes from the liver^42^ (artery, capillary artery, capillary, capillary vein, vein), and performed the analysis using the parameters k = 3 and number of PC = 8.

### Perturbations of zonation in disease states

Monocle2 was used to derive a common zonation trajectory across biological conditions within each cell type. This enabled us to compare the conditions along the trajectory and detect large scale gene expression changes indicative of differential progression. Differentially expressed genes along the trajectory between control (chow diet) and either NAFLD or NASH liver groups, and between the control and CCl_4_-exposed mesenchymal cell groups, were identified using the *conditionTest* function of the tradeSeq package^143^. Significantly perturbed PCGs and lncRNAs (FDR < 0.001) were extracted and used to prepare matched heatmaps for control liver and from diseased liver cells using the *pheatmap* function of Monocle2. In the case of NAFLD and NASH livers, we identified PCGs and lncRNAs where a change in zonation was significant for either NAFLD vs chow diet or for NASH vs. chow diet. We then clustered heatmaps individually from each condition to assign the zonation labels shown in Table S4I.

### Functional enrichment analysis

Sets of PCGs obtained in various analyses were used as input for functional enrichment analysis using DAVID^144^ with default parameters, except that GO FAT terms were used in place of GO DIRECT terms to include a broader range of enrichment terms that were excluded by the default GO DIRECT option. We used a custom script prepared by Dr. Alan Downey-Wall of this laboratory (https://github.com/adowneywall/Davidaggregator) to combine and reformat DAVID output files from multiple gene lists, with each row presenting top enriched annotation clusters and individual columns presenting enrichment score, p-value, FDR and other such data. Top terms from each DAVID annotation cluster with a cluster enrichment score > 3 and top FDR < 0.05 were used in downstream analysis.

### Differential expression analysis between biological conditions

Differential gene expression comparisons between single cell clusters and between biological conditions were performed using the Locally Distinguishing function of the 10x Genomics Loupe Browser (v.6.0). This method implements the negative binomial test based on the sSeq method^141^ with Benjamini-Hochberg correction for multiple tests. Further, the method uses log-normalized average expression values and the distribution of UMIs across the two specific cell populations being compared to calculate fold-change and FDR values for each pair-wise comparison between cell clusters. Genes were identified as showing significant differential expression between control liver and diseased liver (NAFLD, NASH, or CCl_4_-induced liver fibrosis) if they met the thresholds of log2 |fold-change| >1 and FDR <0.05.

### Transition from quiescent HSCs to activated HSCs (myofibroblasts)

Central vein-associated HSCs were extracted from livers of mice treated with CCl_4_ (6 wk treatment) to investigate the transition of gene expression from resting HSCs to collagen-producing myofibroblasts. The central vein-associated HSCs were spatially resolved into resting HSCs and myofibroblasts using Monocle2 (v2.6.419), and the two cell subtypes then characterized using established signature genes^35^. PCGs and lncRNAs showing zonation along the trajectory were identified using generalized linear models and a heatmap was prepared using Monocle2, as described above.

### Construction of gene regulatory networks

Regulatory networks were generated using the R package bigSCale2^96^ for scRNA-seq datasets comprised of all cells that passed the gene expression filters described above. Three separate bigSCale2 networks were generated from scRNA-seq data (GSE166504) from livers of mice fed chow diet (healthy liver) or mice fed HFHFD for either 15 wk (NAFLD liver) or 30 wk (NASH liver). Two other bigSCale2 networks were generated from all qualified mesenchymal cells from the scRNA-seq datasets for control and for CCl_4_-exposed mouse liver (GSE137720) (Table S1G). Cells from each of the 5 datasets were split into individual Seurat objects and then input to bigSCale2. Each of the 5 networks was constructed using default clustering parameters, with granularity set to the highest setting and applying an edge cutoff of the top 99.9 quantile used for correlation coefficient. BigSCale2 retains edges that are expected to represent actual regulatory links by removing genes that did not have a direct edge with a known GO regulator (GO sub-setting step). This was implemented using a list of GO regulators comprised of the union of genes in GO:0006355 “regulation of transcription, DNA-templated” (N=2,776) and genes in GO:000370 “DNA-binding transcription factor activity” (N=1,001). We also included the above list of 52,961 lncRNAs as potential regulators in the GO sub-setting step. Network specifications are shown in Table S5E. The BigSCale2 gene regulatory networks obtained were converted into json files for visualization by Cytoscape^145^ using the *toCytoscape* function of the package iGraph (v1.3.5) (https://igraph.org/r) in R. Specifically, the network json file was imported into Cytoscape using “*Import Network from File”*. Next, we used “*forced- directed”* layout in Cytoscape, where each network’s layout was derived from 10,000 iterations of the Fruchterman-Reingold algorithm^146^ and using the Nogrid parameter (seed=7). The resultant networks and derived subnetworks shown in the various figures are available at https://tinyurl.com/scLiverNetworksKarriWaxman.

### Analysis of gene regulatory networks

Gene modules in each network were discovered using the Glay community cluster algorithm^147^ in Cytoscape, whereby the overall network topology was used to subdivide the networks generated by bigSCale2 into functional modules . Genes in each module were input to DAVID^144^ for gene ontology enrichment analysis. Node rank was calculated for each of four key bigSCale2 network metrics (Betweenness, Closeness, Degree, PAGERANK). PCGs that ranked within the top 100 nodes for any one of the above four network metrics were deemed to be network-essential nodes that correspond to regulatory PCGs; and lncRNAs that ranked within the top 50 nodes for any one of the above four network metrics were deemed to be network-essential nodes identifying regulatory lncRNAs (Table S5C).

### Master regulators

We extracted subnetworks comprised of the network-essential regulatory PCGs and regulatory lncRNAs defined above, i.e., all top-100 ranked PCG nodes and all top-50 ranked lncRNA nodes. We then recalculated the network metrics to identify critical nodes that are deemed network-essential regulators with regards to expression of the original set of regulators; these were designated master regulators. For this analysis, we ranked the nodes of the extracted networks using 5 different network metrics: Stress, Degree, Betweenness, Closeness, and Neighborhood Connectivity, as calculated using the “*Analyze Network*” option in Cytoscape. We then designated as master regulators the top 5 ranked nodes for each of the 5 metrics, which were considered independently in the ranking (Table S5C, columns AP-BU). We excluded from the master regulator analysis the PageRank centrality metric (used in the original ranking, above), as it tended to select nodes with low Degree ranking due to the high degree of disconnectedness in these master regulator networks.

### IPA Analysis

We used Qiagen’s Ingenuity Pathway Analysis (IPA) software (https://digitalinsights.qiagen.com) to validate the regulatory role of a total of the 20 master regulators derived from all 5 bigSCale2 networks that were identified as DNA-binding proteins. Putative gene targets of these 20 DNA-binding master regulators were provided as input to IPA. IPA’s core analysis of these input gene lists was used to identify Upstream Regulators, along with the top enriched canonical pathways, diseases, and toxicological functions. The resulting IPA output for all 20 master regulators is included in Table S5G.

### Enrichment heatmaps

PCGs that made direct connections to a network-essential regulatory lncRNA in any of the bigSCale2 networks were used as input to Metascape with default parameters^119^. To identify pathways that are shared between, or are specific, to a lncRNA’s gene targets, we performed multi-list comparative analysis within and across the networks. The resulting clustered heatmaps depict -log10 P-values of the top enriched pathways across multiple lncRNAs target lists.

### Triplex Domain Finder analysis

Regulatory lncRNAs identified for each gene network using bigSCale2 metrics (see above) were analyzed using the algorithm Triplex Domain Finder (TDF)^120^ to determine their capacity to form lncRNA-genomic DNA triplexes with the promoter sequences of their target genes. Target genes were defined as all PCGs that make a direct connection to the lncRNA in the corresponding bigSCale2 network. A background gene set was defined as the set of PCGs expressed in at least 0.5% of the cells in the network, excluding all genes that were part of a bigSCale2 network derived from the same cell population used to identify the regulatory lncRNA being evaluated for triplex formation, but constructed using an edge cutoff at the 95% quantile (instead of the 99.9% quantile) correlation coefficient threshold. This relaxed threshold was implemented to obtain a bigSCale2 network specifically for the purpose of background gene identification, and with the goal of removing from the background gene set any gene that might be a target of the network-essential regulatory lncRNA when using a less stringent correlation threshold of >95% quantile. Fisher exact test was used to assess the significance (at FDR <0.05) of the observed number of target gene promoters predicted by Triplex Domain Finder to form lncRNA-DNA triplexes, compared to the number of background gene set promoters predicted to form triplexes with the same lncRNA. The Triplex Domain Finder-predicted gene binding targets of each regulatory lncRNA were used to derive lncRNA─gene directed subnetworks for each condition (Fig. S15, Fig. S16) using Cytoscape^145^, and are available at https://tinyurl.com/scLiverNetworksKarriWaxman.

## Supplemental information

### Supplementary figure titles and legends

**Fig. S1.**
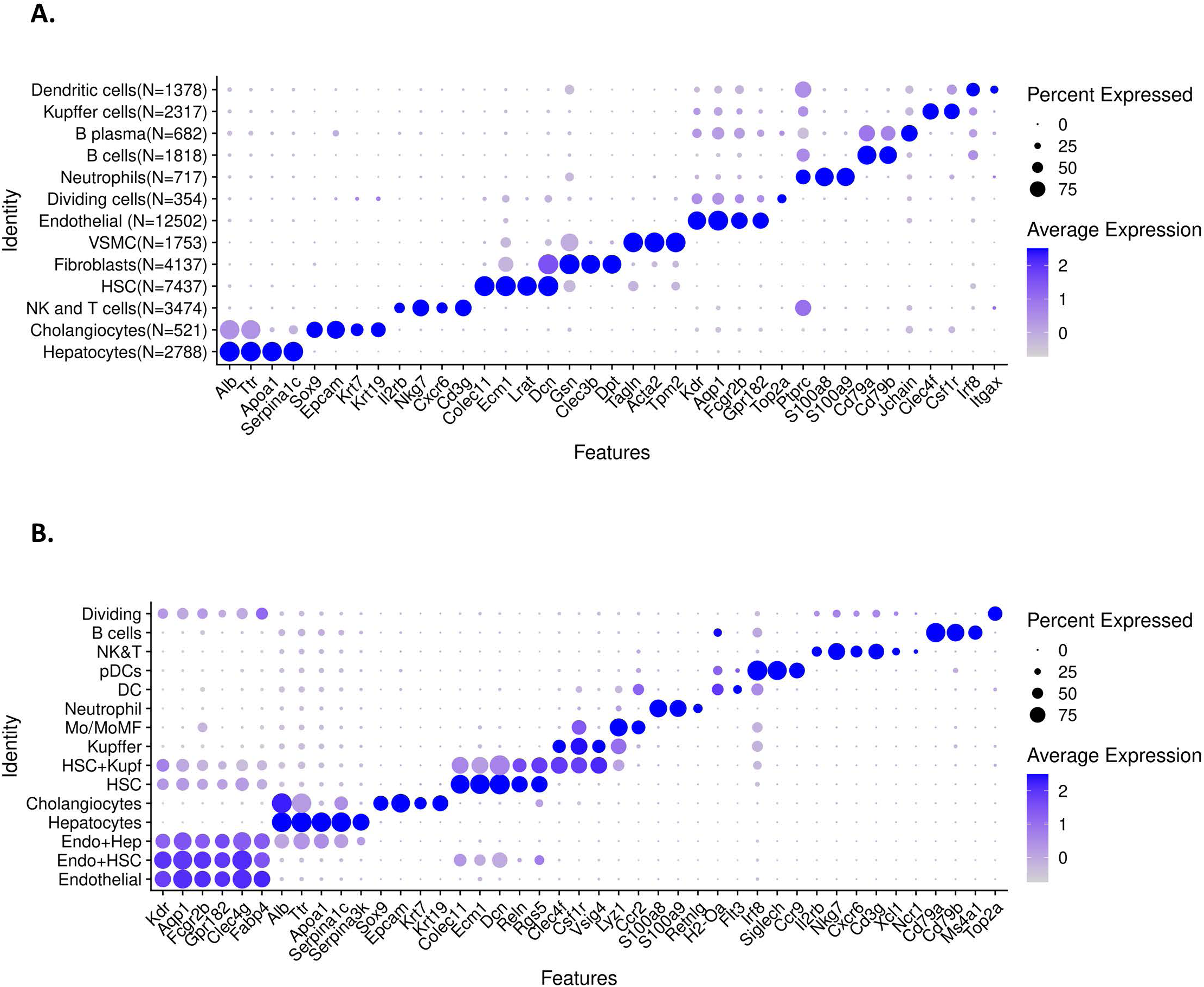
Identification of liver cell clusters from healthy mouse liver. **A.** Dot plot showing average expression values for marker genes (shown along X-axis) across the 13 hepatic cell clusters (Y-axis) shown in the UMAP in Fig. 2A. Those cell clusters were based on hepatocytes and NPCs from four datasets from control mouse livers. **B.** Dot plot showing average expression values for marker genes (shown along X-axis) across the 12 hepatic cell clusters (Y-axis) shown in the UMAP in Fig. 4A. Those cell clusters were based on liver cells isolated from healthy mice (chow-fed diet), and from mice fed HFHFD for 15 wk to induce simple steatosis (NAFLD), or for 30-34 wk to induce NASH.

**Fig. S2.**
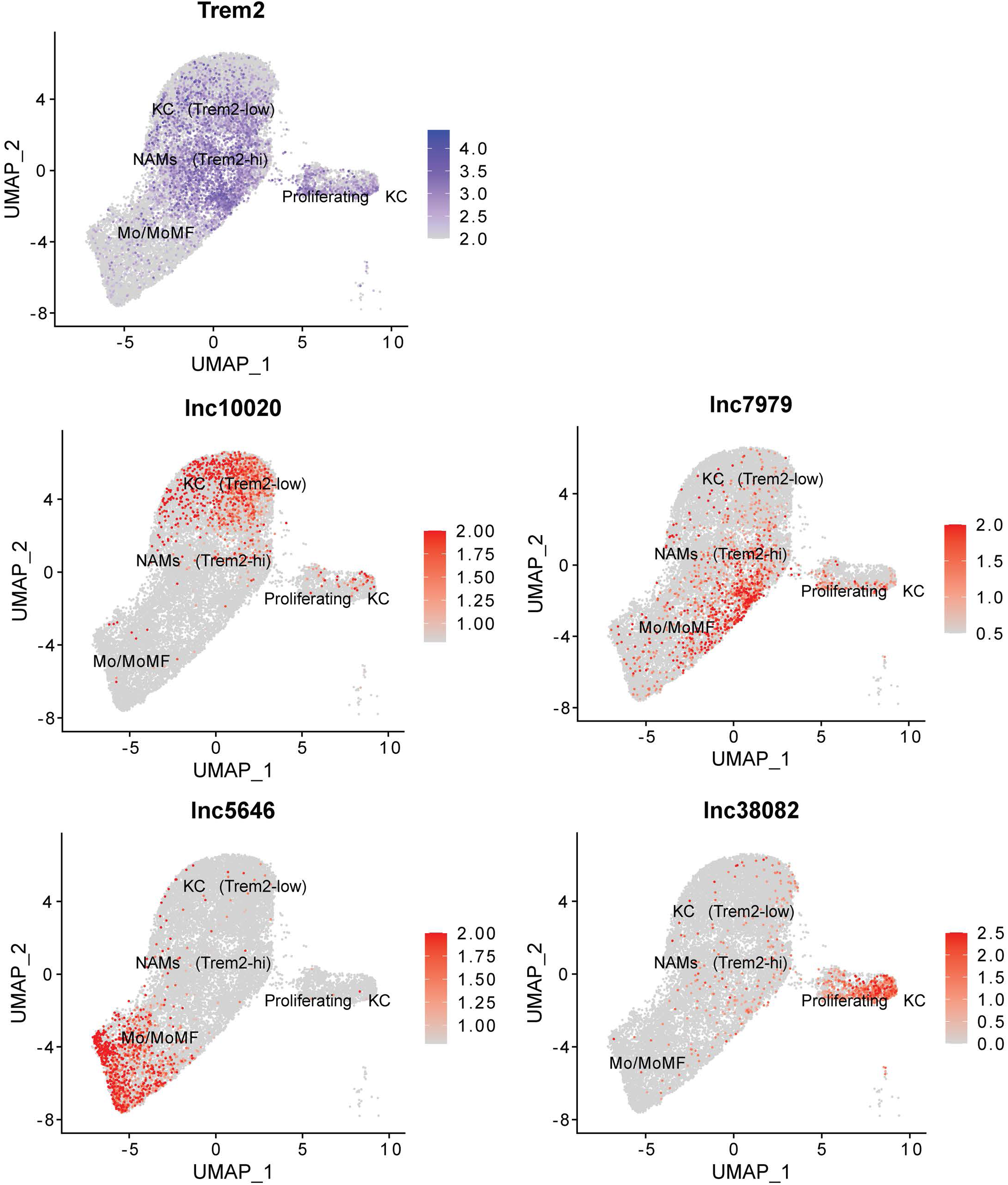
Kupffer cell subpopulations. Feature plots presenting marker gene expression for Trem2, and for 4 lncRNAs that are markers for different Kupffer subpopulations. UMAP is as shown in Fig. 4G.

**Fig. S3.**
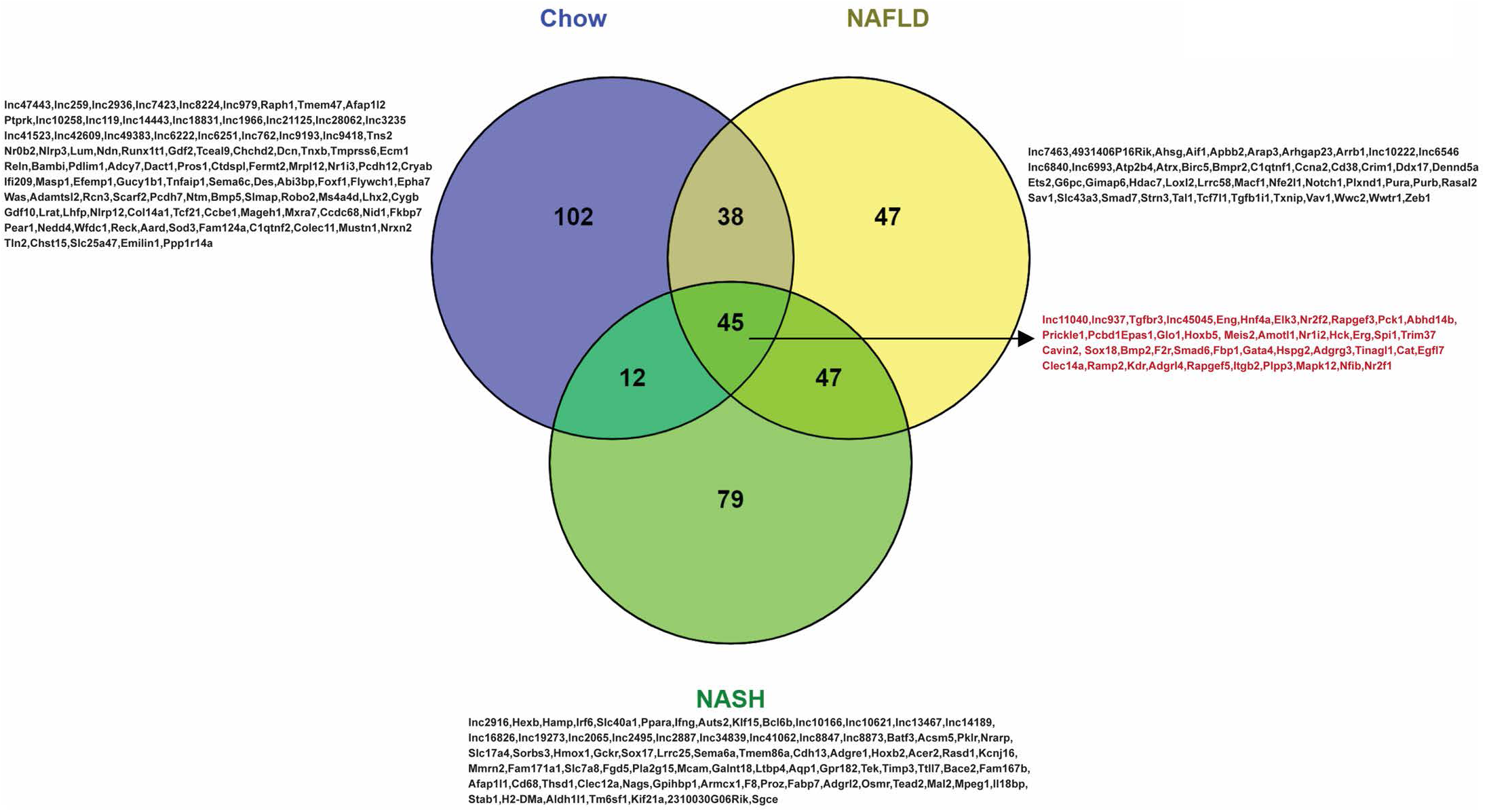
Shared network-essential regulators in healthy and diseased (NAFLD and NASH) mouse livers. Venn diagram showing overlap between network-essential PCGs and lncRNAs from control, NAFLD and NASH gene regulatory networks from Fig. 6 and Table S5.

**Fig. S4.**
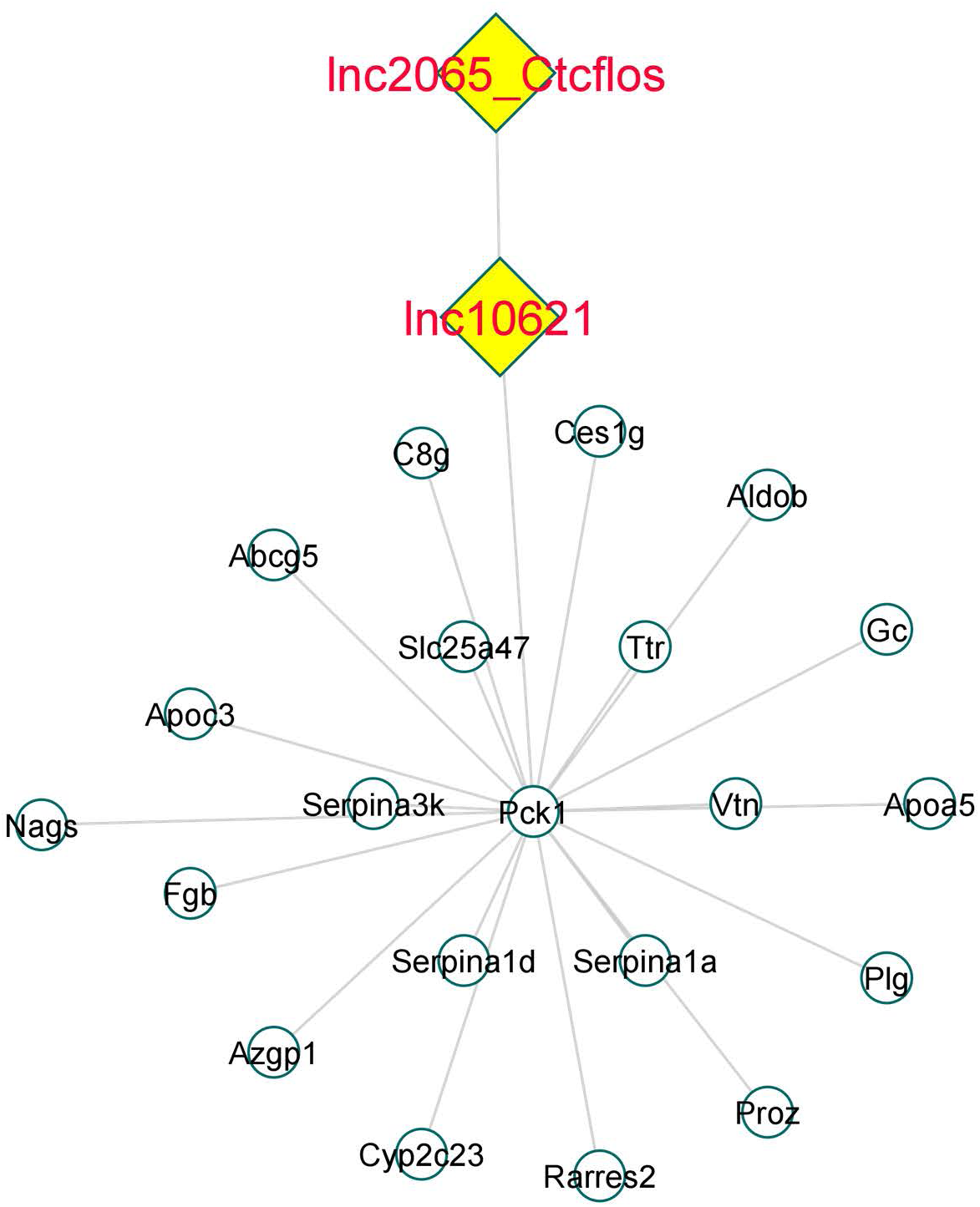
Ctcflos regulation of Pck1. Subnetwork showing Ctcflos (lnc2065), an essential network regulatory gene in the NASH network, which makes connections with Pck1 via lnc10621.

**Fig. S5.**
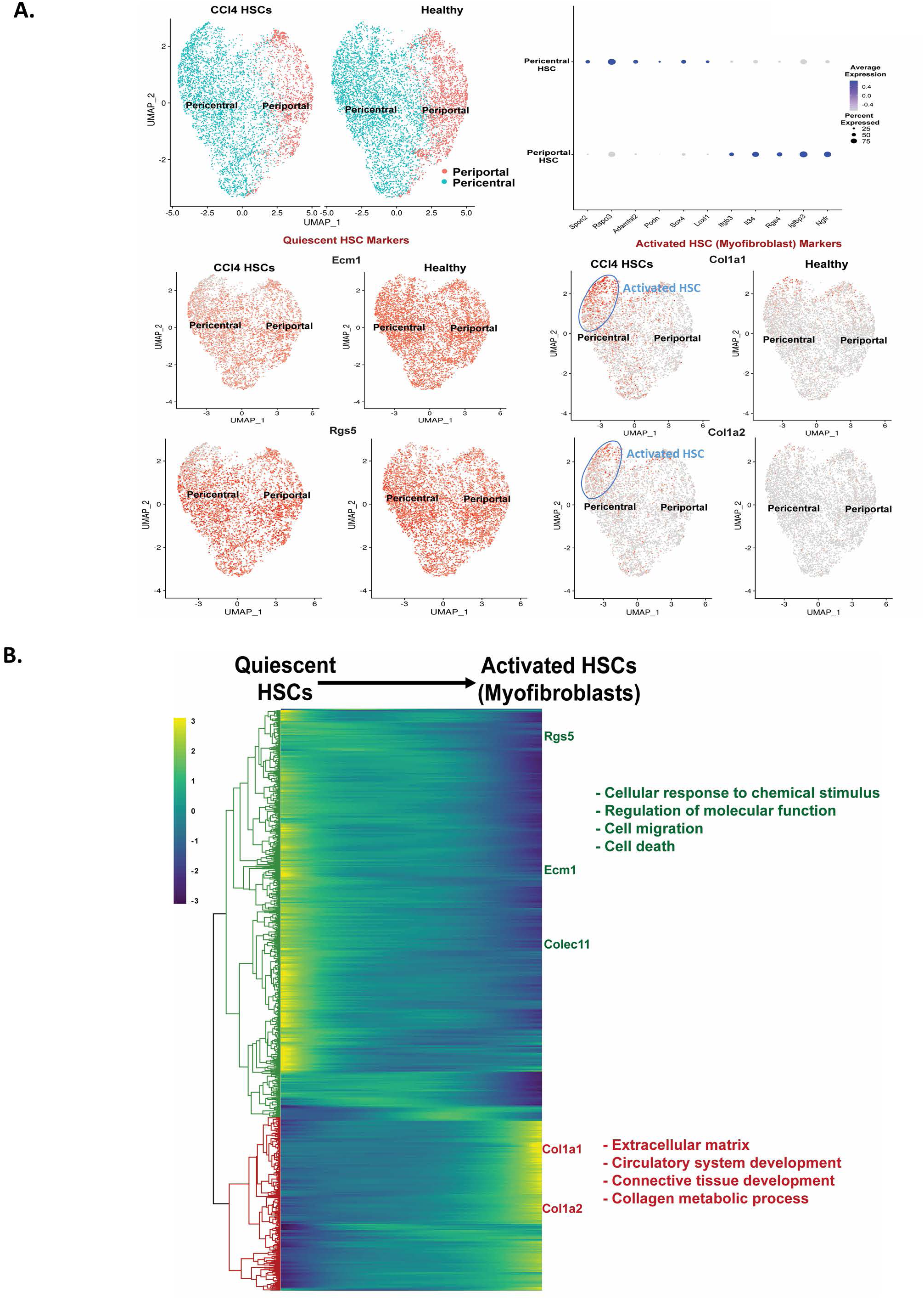
Differentiation of quiescent HSCs into myofibroblasts. **A.** UMAP showing HSCs zonation clusters (top *left*) and corresponding DotPlot of known HSC marker genes (top *right*). Shown are feature plots for genes that are markers for quiescent HSCs (Ecm1, Rgs5) and for activated HSCs (myofibroblast) (Col1a1, Col1a2). **B.** Heatmap showing relative expression patterns for PCGs that are differentially expressed between quiescent pericentral HSCs and myofibroblasts from CCl_4_-exposed mouse liver mesenchymal cells, along with their top functional enrichment terms, listed on the right. See Fig. 7E for corresponding heat map for lncRNA genes.

**Fig. S6.**
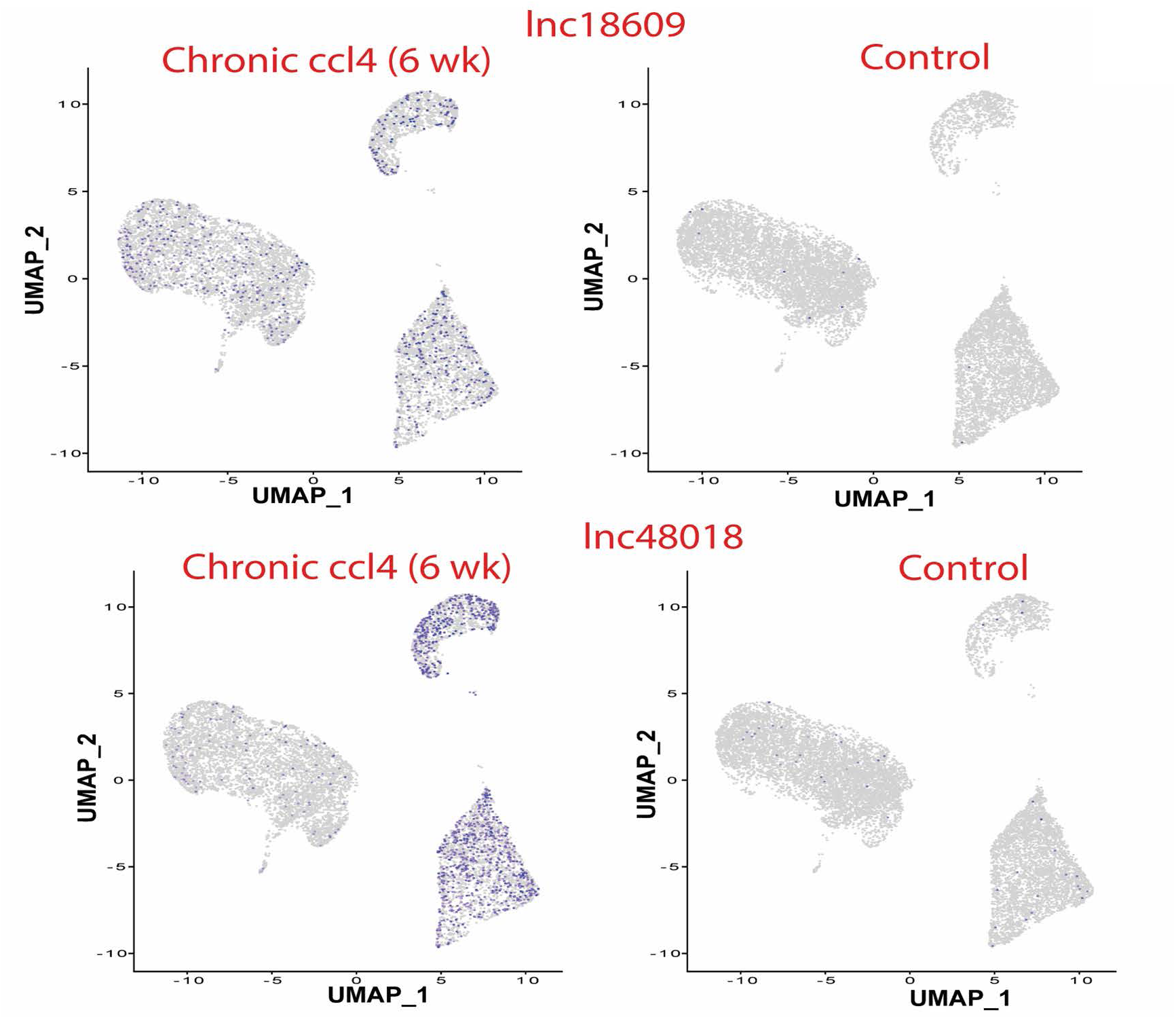
Feature plots showing examples of lncRNAs that are differentially expressed across multiple mesenchymal cell subpopulations.

**Fig. S7.**
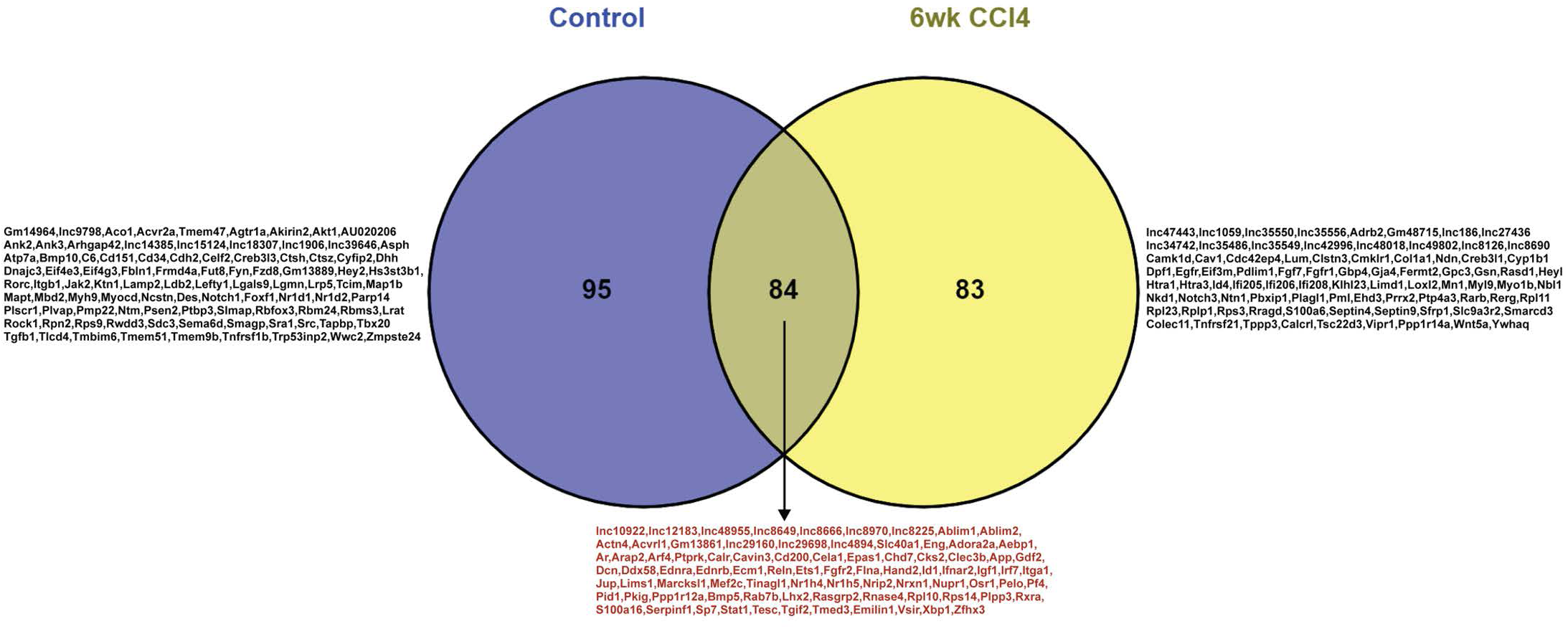
Shared gene regulatory network regulators in healthy and CCl_4_-induced fibrotic liver. Venn diagram showing overlap between network-essential PCGs and lncRNAs from the healthy and fibrotic liver gene regulatory networks shown in Fig. 8.See details in Table S5.

**Fig. S8.**
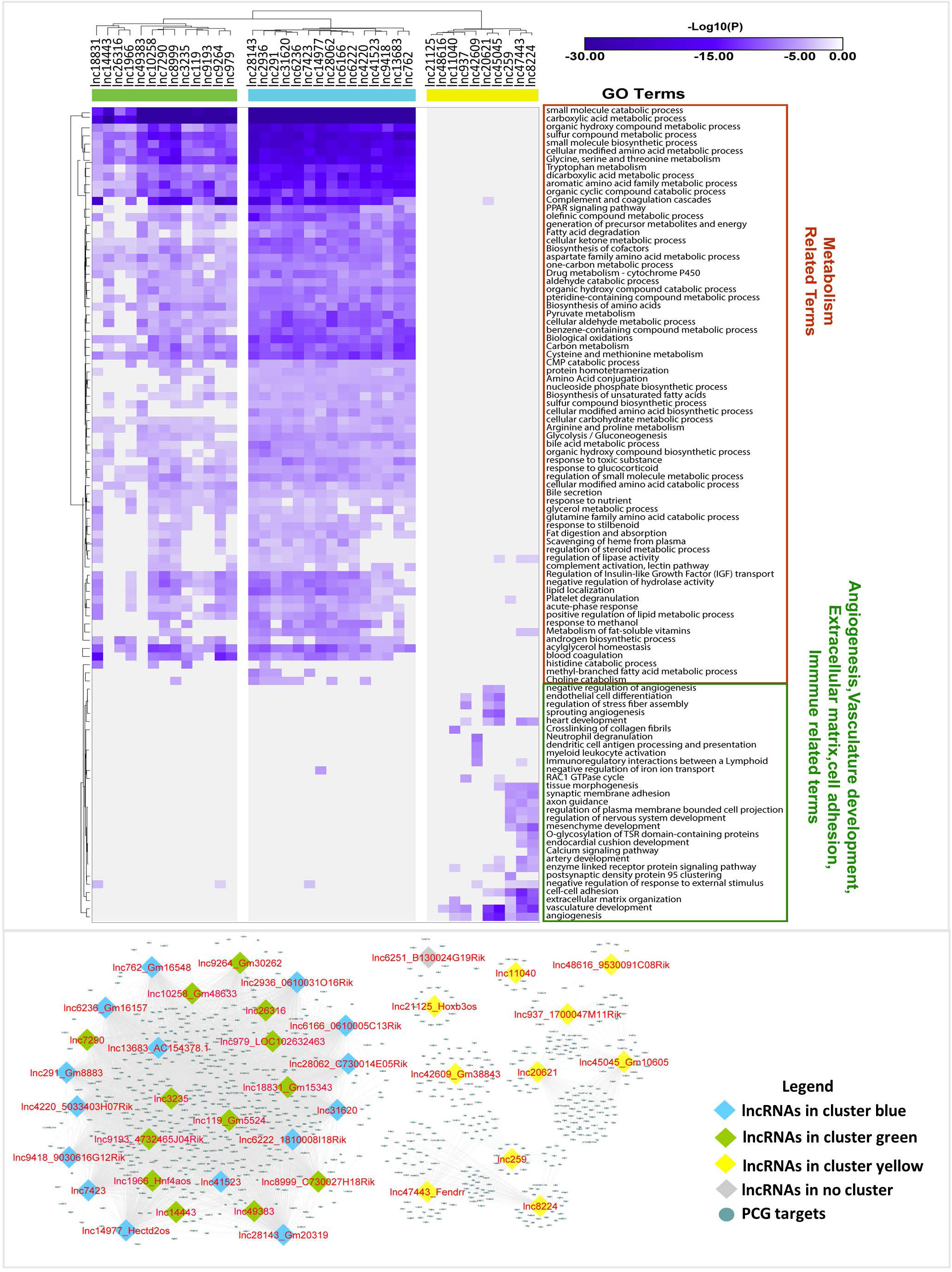
Clustering of 39 regulatory lncRNAs from chow-fed (control) liver gene regulatory network. **Top**: PCG targets that make a direct connection with regulatory lncRNAs in the chow fed liver network of Fig. 6A were provided as input to Metascape to obtain top functional enrichment terms based on -log10 (P-value). The color scale represents -log10(P) values, with darker purple coloration indicating more significant p-values. Two-way hierarchical clustering of the functional enrichment heatmap split the 39 lncRNAs into 3 clusters (columns colors) based on their patterns of shared functions. Target genes of lncRNAs from two of the major functional clusters (green, blue) were variously enriched for diverse metabolic processes, and a third (yellow) was functionally enriched for angiogenesis and vasculature development, extracellular matrix, and cell-cell adhesion, as shown on the right. **Bottom**: subnetwork of the network shown in Fig. 6A. This subnetwork is comprised of the network-essential 39 regulatory lncRNAs and their target genes, which clustered to give gene modules that mirror the clustering results obtained for the same lncRNAs based on their functional group enrichments, shown on top. The subnetwork was derived from the network in Fig. 6A by removing all PCG regulators and their gene targets, and by retaining regulatory lncRNAs and their direct protein coding genes connections, which were used as input for the functional enrichment analysis shown at the top. The regulatory lncRNAs (nodes) are color coded to match the lncRNA cluster colors shown in the heat map. Gene targets for lnc6251 (bottom, gray node) did not show enrichment for any functional terms and was therefore excluded from the enrichment heatmap (top).

**Fig. S9.**
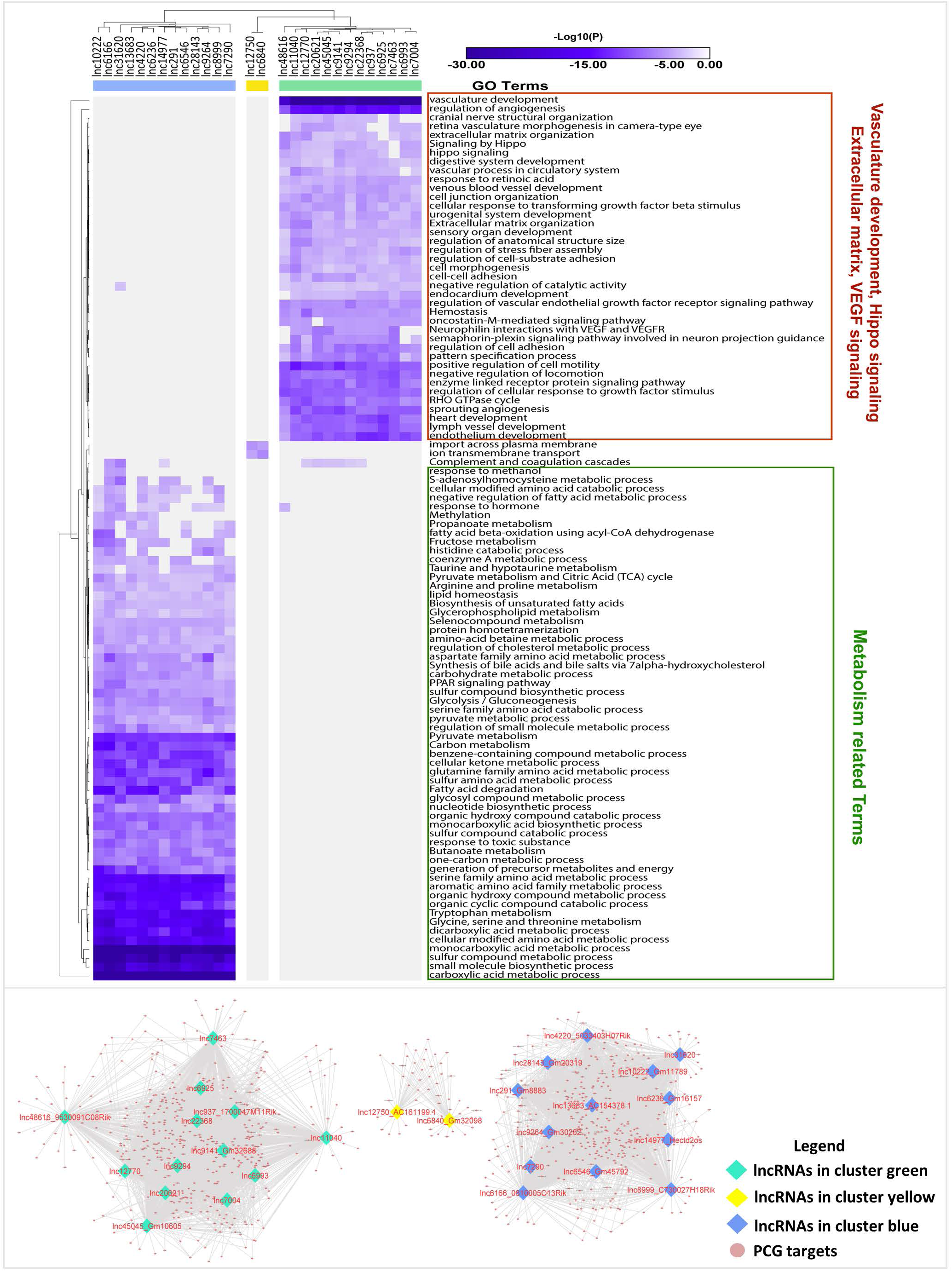
Clustering of 28 regulatory lncRNAs from NAFLD gene regulatory network. **Top**: Analysis of network- essential regulatory lncRNAs from NAFLD network of Fig. 6B, as described in Fig. S8. Two-way hierarchical clustering identified 3 clusters of regulatory lncRNAs, with target genes of lncRNAs from cluster blue enriched for carboxylic acid and other metabolic processes, cluster green lncRNA targets enriched for vasculature development, angiogenesis, hippo signaling, extracellular matrix and cell motility, and the target genes of lncRNAs from cluster yellow, which is comprised of only 2 lncRNAs (lnc6840, lnc12750), specifically enriched for import across plasma membrane and ion transport. **Bottom**: subnetwork of the network shown in Fig. 6B. It is comprised of the 28 regulatory lncRNAs and their target genes (excluding regulatory PCGs) and was obtained as described in Fig. S8. The 28 regulatory lncRNAs clustered to give gene modules that mirror the clustering results obtained for the same lncRNAs based on their functional group enrichments, shown on top.

**Fig. S10.**
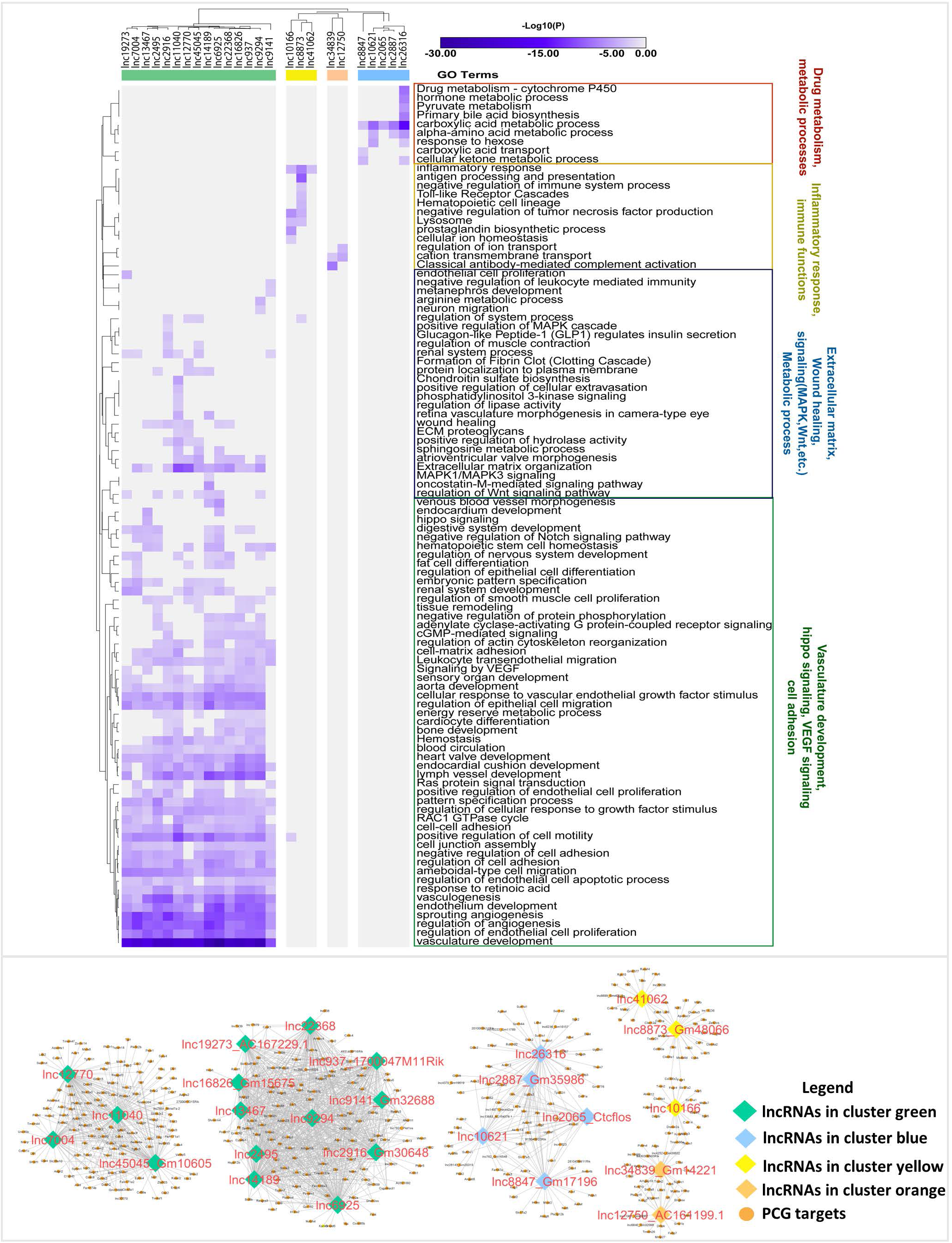
Clustering of 25 regulatory lncRNAs from NASH gene regulatory network. **Top**: Analysis of network- essential regulatory lncRNAs from NASH network of Fig. 6C, as described in Fig. S8. Two-way hierarchical clustering identified 4 clusters of regulatory lncRNAs, with the target genes of lncRNAs from cluster green enriched for vasculature development, angiogenesis, and cell motility, and with a subset of lncRNAs in this cluster enriched for extracellular matrix, wound healing, and regulation of Wnt signaling. The target genes of lncRNAs from cluster yellow (n=3 lncRNAs) were enriched for inflammatory response gene targets, while the target genes of lncRNAs from cluster orange (n=2 lncRNAs) were enriched for cation transmembrane transport. The targets of cluster blue lncRNAs (n=5) were enriched for carboxylic acid and other metabolic processes. **Bottom**: subnetwork of the network shown in Fig. 6C. It is comprised of the 25 regulatory lncRNAs and their target genes (excluding regulatory PCGs) and was obtained as described in Fig. S8. The 25 regulatory lncRNAs clustered to give gene modules that mirror the clustering results obtained for the same lncRNAs based on their functional group enrichments, shown on top.

**Fig. S11.**
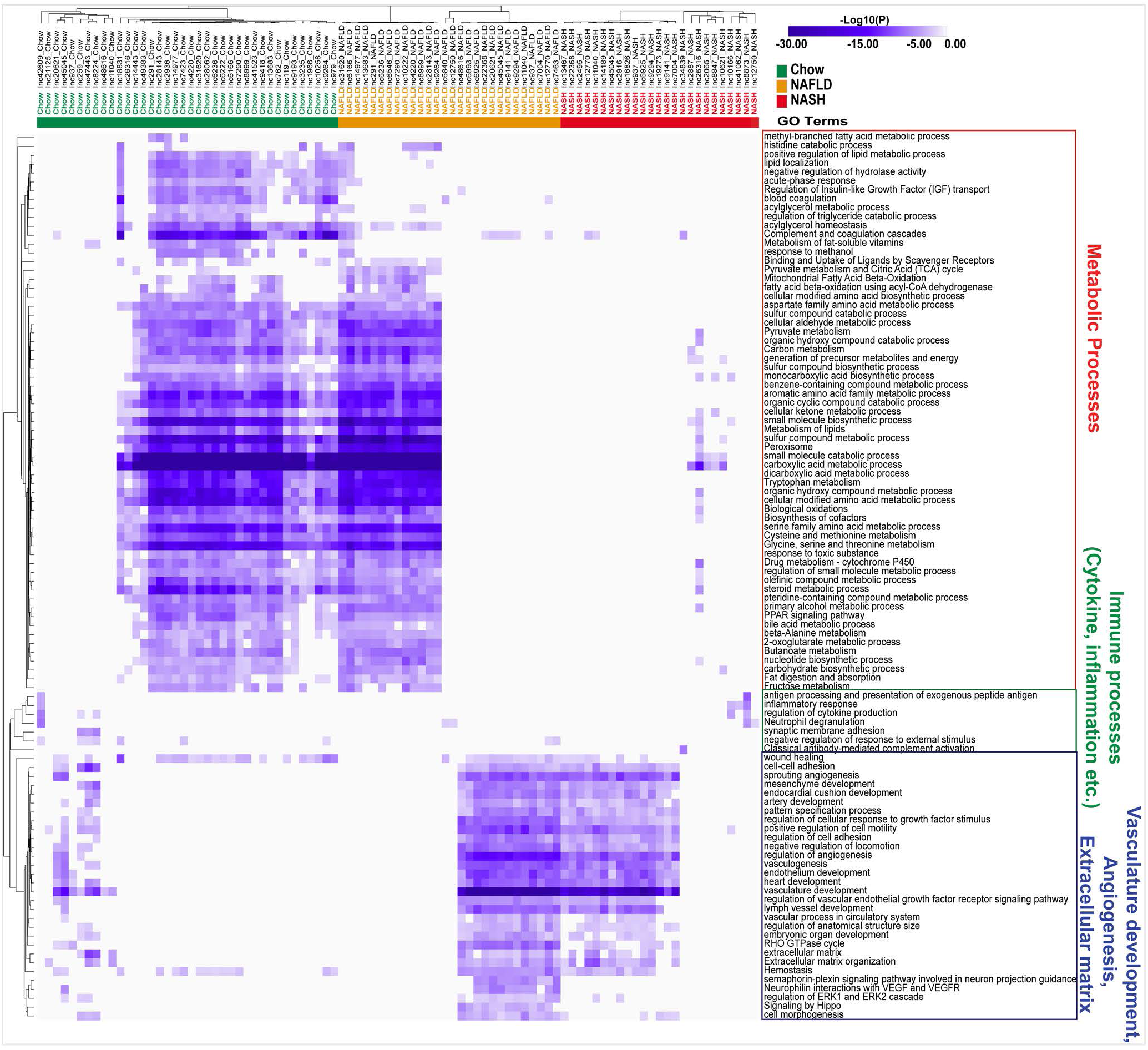
Combined clustering of 92 regulatory lncRNAs from Chow, NAFLD and NASH liver gene regulatory networks. Shown is a merged functional enrichment heatmap showing the top enriched terms (as rows) for genes targets of all network-essential regulatory lncRNAs (as columns) from the chow, NAFLD, and NASH networks combined and analyzed as described in Fig. S8. The NAFLD and NASH network-derived regulatory lncRNAs showed more extensive and more significant enrichment for vasculature development and angiogenic processes than those from control liver, whereas the NAFLD and control network regulatory lncRNAs both showed enrichment for diverse metabolic processes. Many fewer regulatory lncRNAs showed target gene enrichment for immune-related processes. Overall, this figure compares regulatory lncRNAs and their target functional terms across biological conditions.

**Fig. S12.**
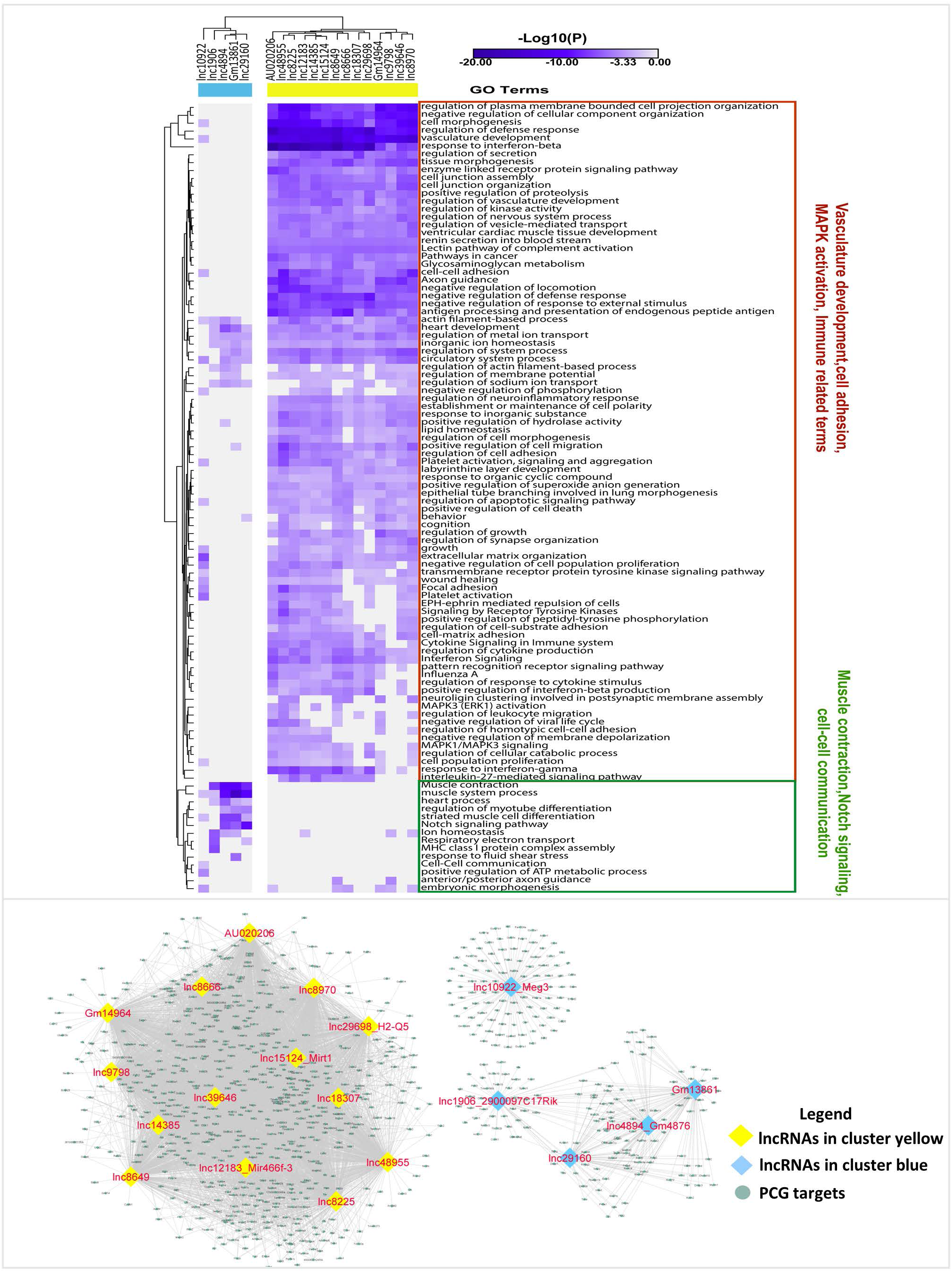
Clustering of 19 regulatory lncRNAs from healthy liver mesenchymal cell gene regulatory network. **Top**: Analysis of network-essential regulatory lncRNAs from the healthy liver mesenchymal network of Fig. 8A, as described in Fig. S8. Two-way hierarchical clustering identified 2 clusters of regulatory lncRNAs, with the target genes of lncRNAs from cluster blue (n=5 lncRNAs) significantly enriched for terms related to muscle contraction, notch signaling, and ion homeostasis, and those of cluster yellow (n=14 lncRNAs) strongly enriched for diverse biological processes, including cell morphogenesis, regulation of defense response, vascular development, and interferon responses. **Bottom**: subnetwork of the network shown in Fig. 8A. It is comprised of the 19 regulatory lncRNAs and their target genes (excluding regulatory PCGs) and was obtained as described in Fig. S8. The 19 regulatory lncRNAs clustered to give gene modules that mirror the clustering results obtained for the same lncRNAs based on their functional group enrichments, shown on top.

**Fig. S13.**
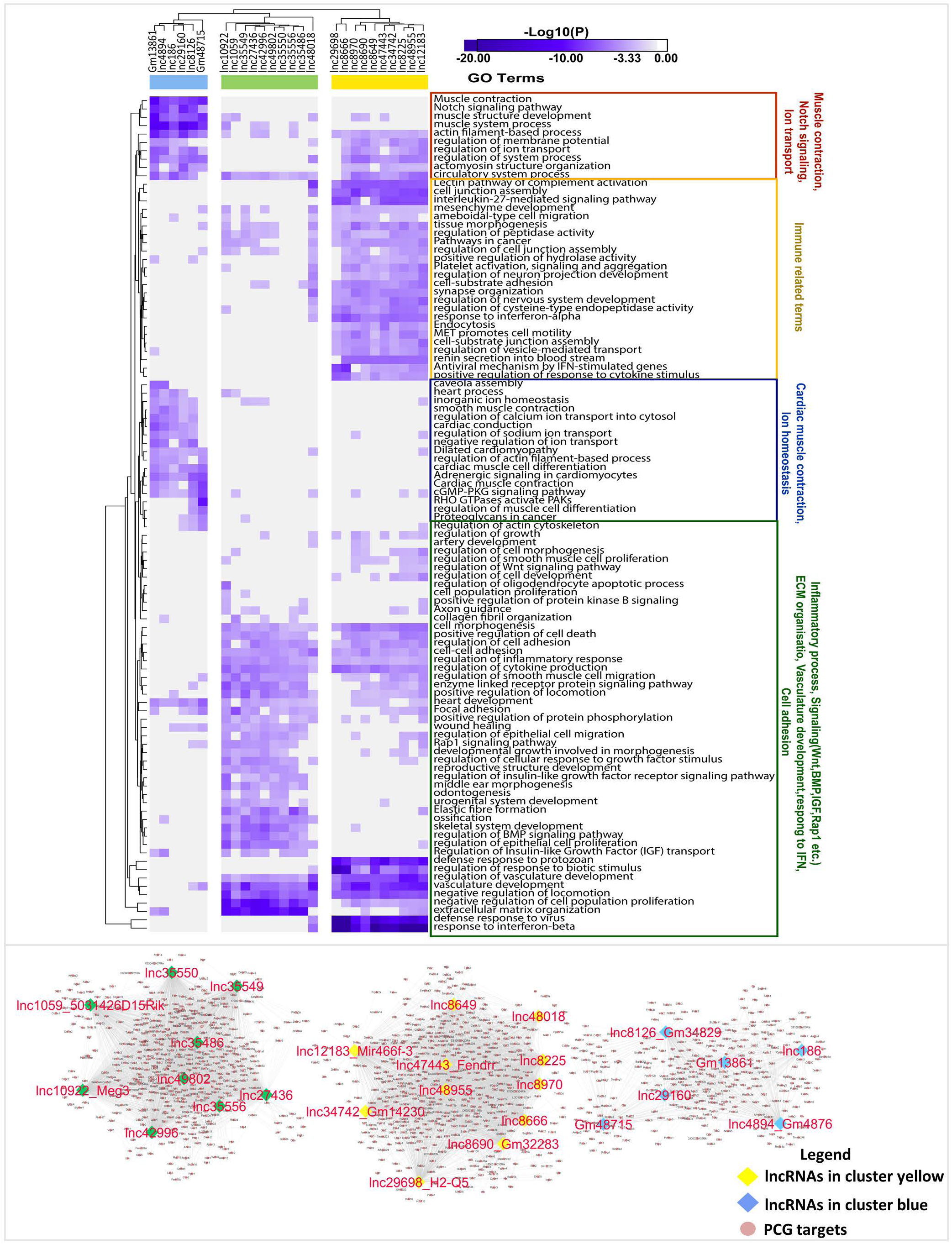
Clustering of 26 regulatory lncRNAs from CCl_4_-induced liver fibrosis gene regulatory network. **Top**: Analysis of network-essential regulatory lncRNAs from the CCl_4_-treated mesenchymal cell network of Fig. 8B, as described in Fig. S8. Two-way hierarchical clustering identified 3 clusters of regulatory lncRNAs (blue, green, yellow), with the target genes of lncRNAs from these clusters respectively showing strongest enrichments and greatest specificities for: muscle contraction and notch signaling and related processes (blue); extracellular matrix organization and other terms (green); and cell junction assembly, response to interferon-beta, and other terms (yellow). **Bottom**: subnetwork of the network shown in Fig. 8B. It is comprised of the 26 regulatory lncRNAs and their target genes (excluding regulatory PCGs) and was obtained as described in Fig. S8. The 26 regulatory lncRNAs clustered to give gene modules that mirror the clustering results obtained for the same lncRNAs based on their functional group enrichments, shown on top.

**Fig. S14.**
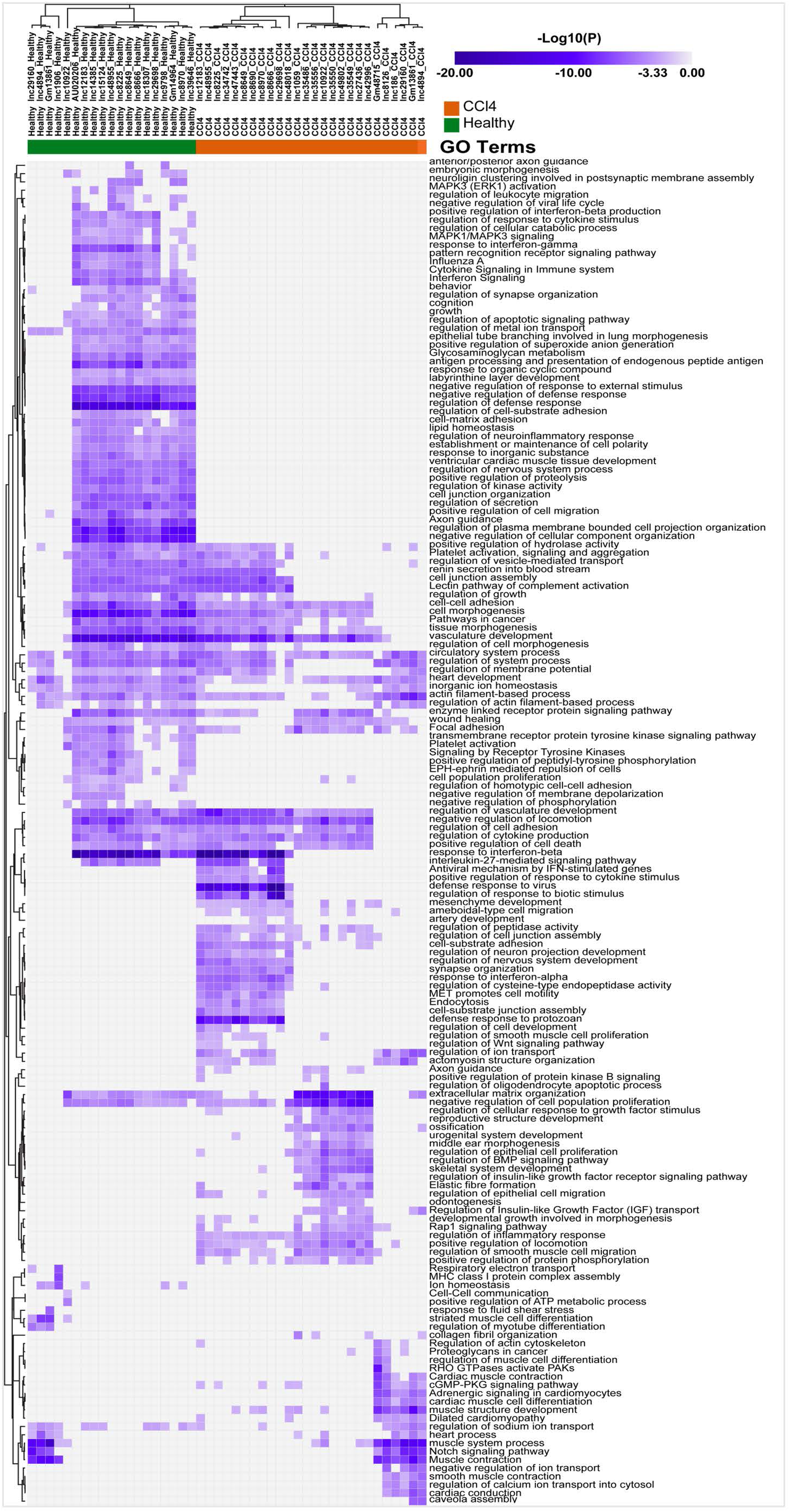
Combined clustering of 45 network-essential regulatory lncRNAs from Healthy and CCl_4_-induced liver fibrosis gene regulatory networks. Shown is a merged functional enrichment heatmap showing the top enriched terms (as rows) for genes targets of all network-essential regulatory lncRNAs (as columns) from the healthy and fibrotic liver mesenchymal gene regulatory networks combined and analyzed as described in Fig. S8. Shown are pathways that were either common or specific to control versus CCl_4_-treated mesenchymal lncRNA gene targets. Overall, this figure compares regulatory lncRNAs and their target functional terms across two biological conditions.

**Fig. S15.**
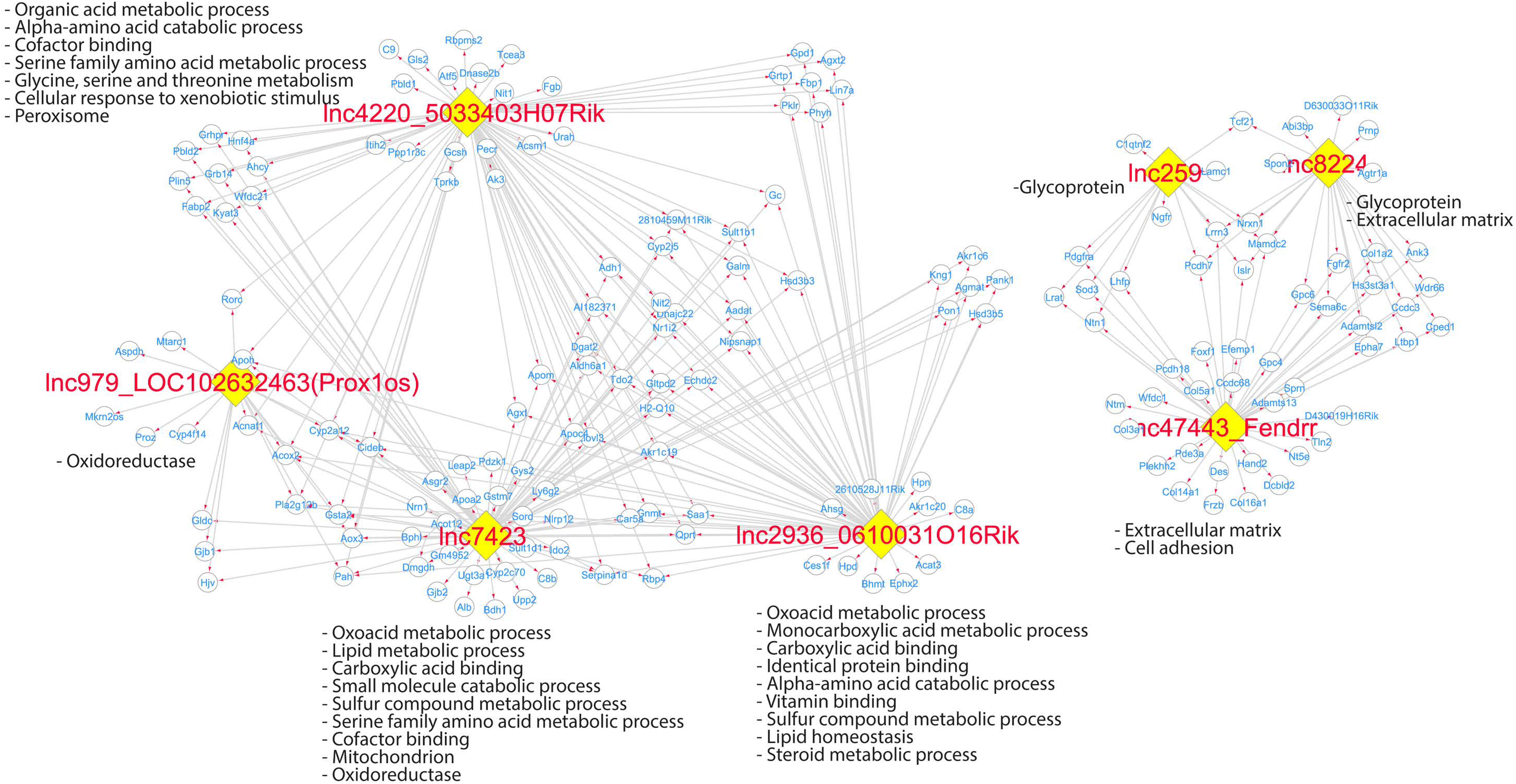
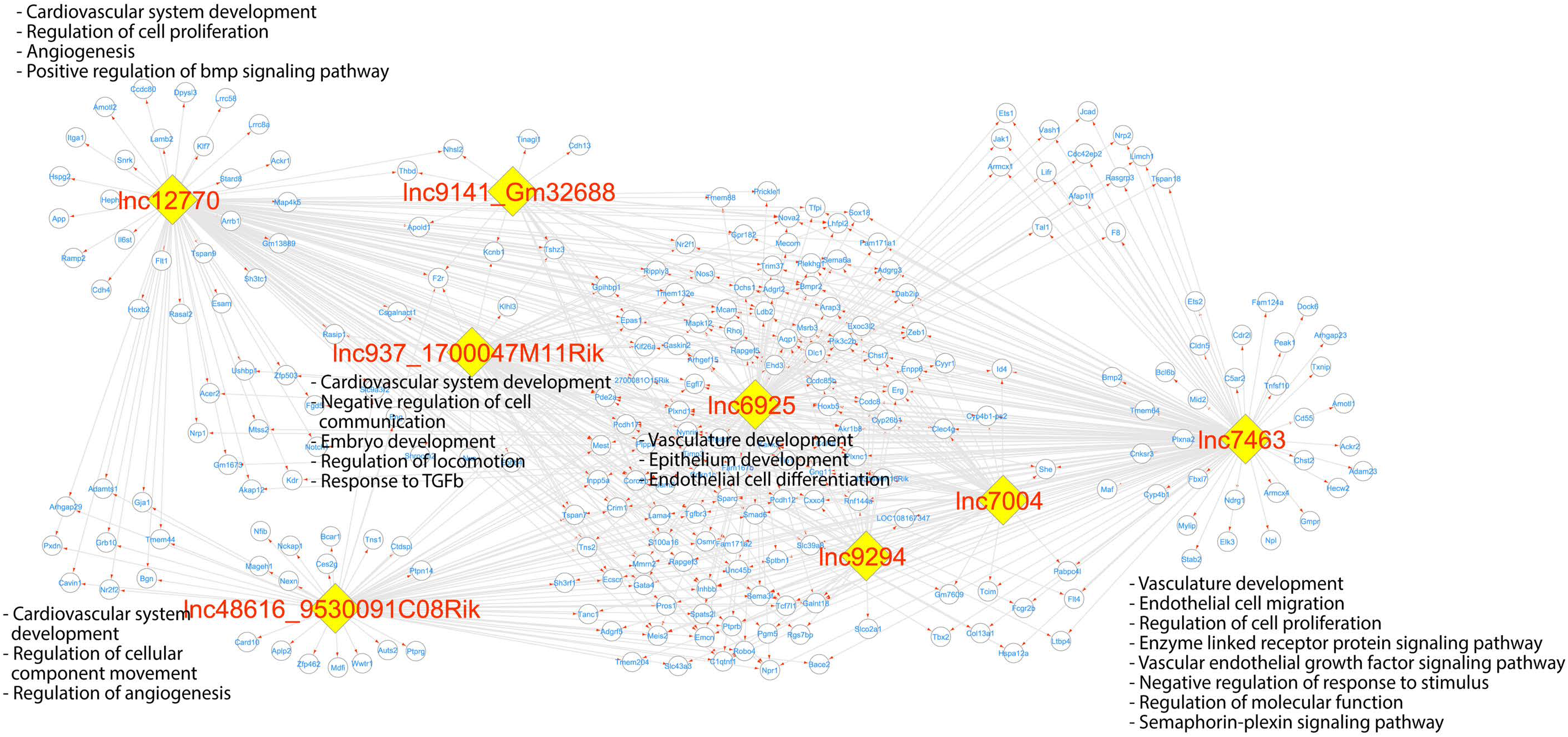
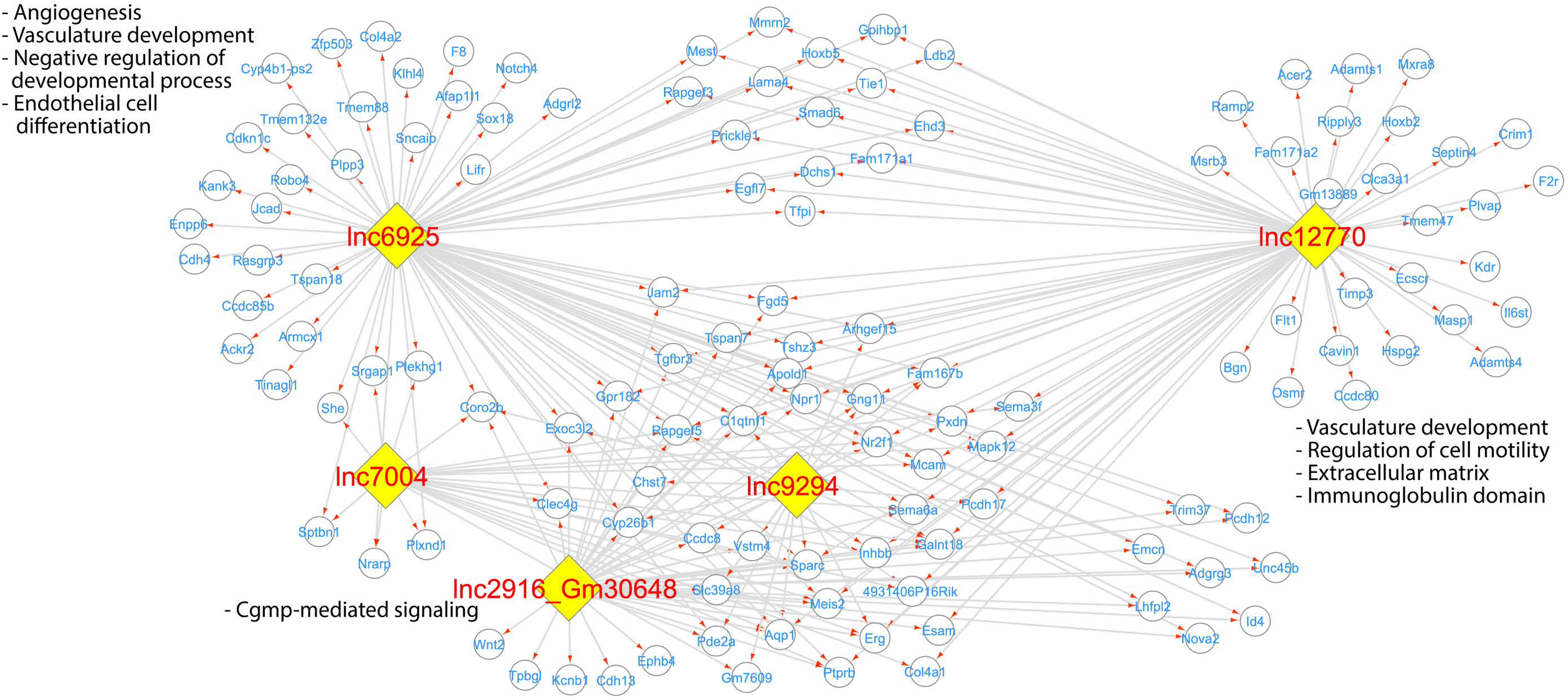
LncRNA-PCG regulatory network based on genes predicted to be involved in lncRNA-genomic DNA triplex binding. Shown are regulatory subnetworks for those lncRNAs that are enriched for forming triplex interactions (based on TDF analysis) with their protein coding gene (PCG) targets identified using bigSCale2 gene regulatory networks from Chow diet (control) liver (**A**), NAFLD liver (**B**), and NASH liver (**C**) (see Fig. 6). The networks shown here are subsets of those shown at the bottom of Fig. S12 and Fig. S13, respectively, insofar as only the regulatory lncRNAs and their protein-coding gene targets that form triplexes based on TDF are shown.

**Fig. S16.**
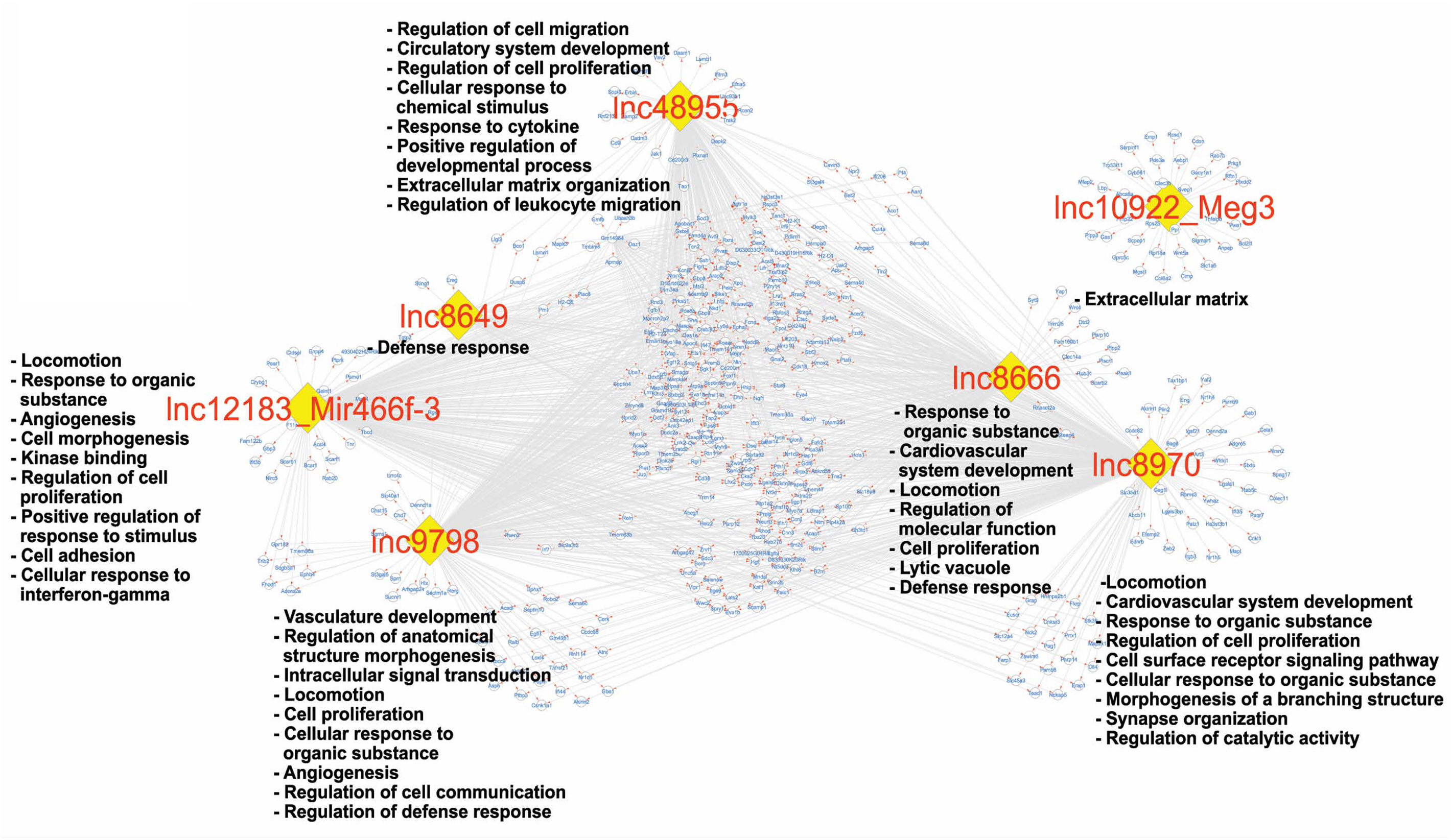
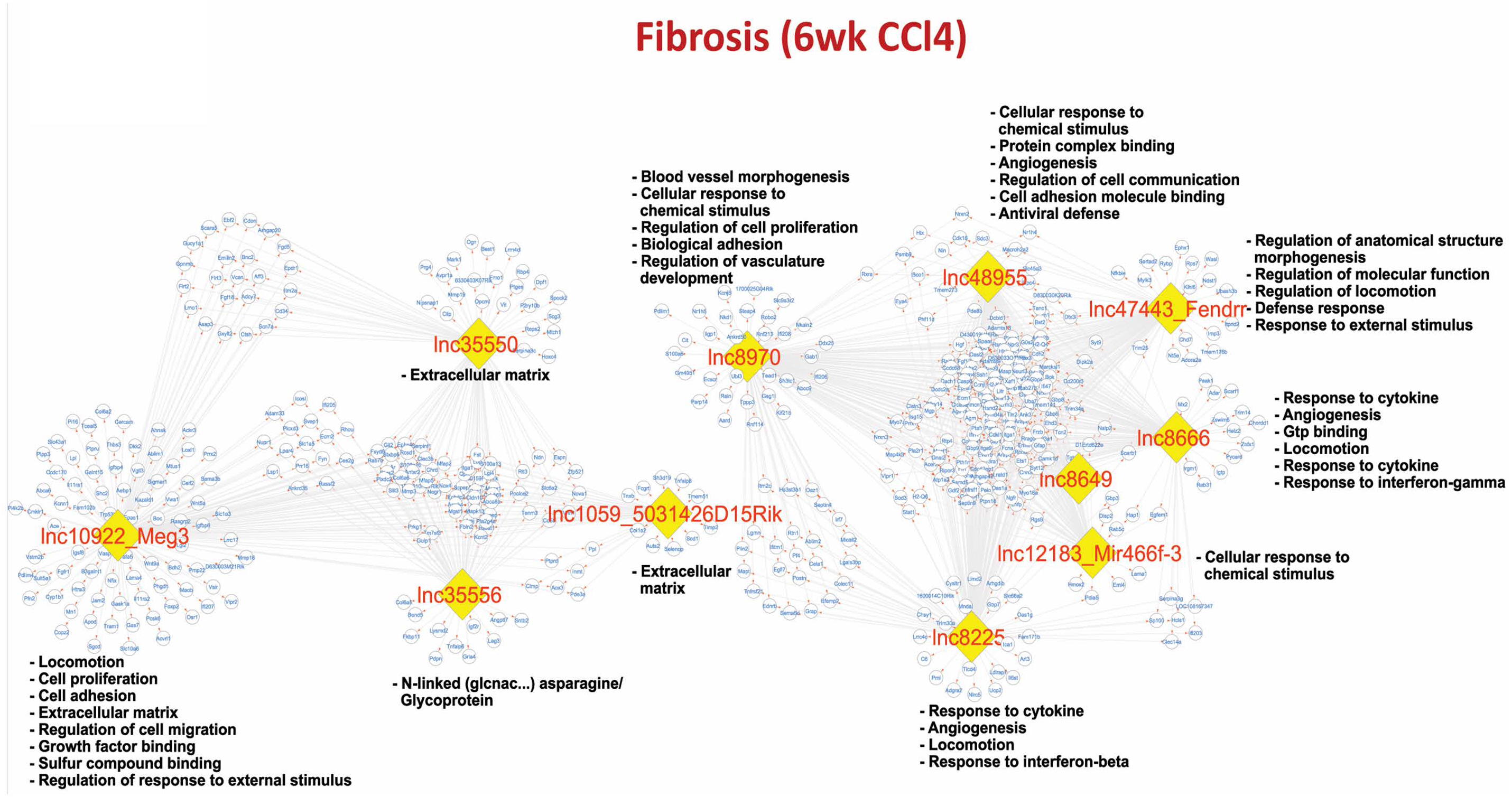
LncRNA-PCG regulatory network based on genes predicted to be involved in lncRNA-genomic DNA triplex binding. Shown are regulatory subnetworks for those lncRNAs that are enriched for forming triplex interactions (based on TDF analysis) with their PCG targets identified using bigSCale2 gene regulatory networks for mesenchymal cells from control liver (**A**) and from CCl_4_-induced fibrotic liver (**B**) (see Fig. 8). The networks shown here are subsets of those shown at the bottom of Figs. S8, S9 and S10, respectively, insofar as only the regulatory lncRNAs and their protein-coding gene targets that form triplexes based on TDF are shown. **Example**: In Fig. 8A, the lncRNA Meg3 is connected to many PCGs and lncRNAs, some of which have their own connections. In Fig. S12, where only regulatory lncRNAs and their direct connections with other regulatory lncRNAs or protein coding genes are shown, Meg3 is in an isolated subnetwork comprised only of its direct target genes. Finally, in Fig. S16A, the subnetwork with Meg3 is also in an isolated subnetwork, one that has even fewer genes than in Fig. S12, because only the targets that are predicted to form a triplex with Meg3 are shown.

**Fig. S17.**
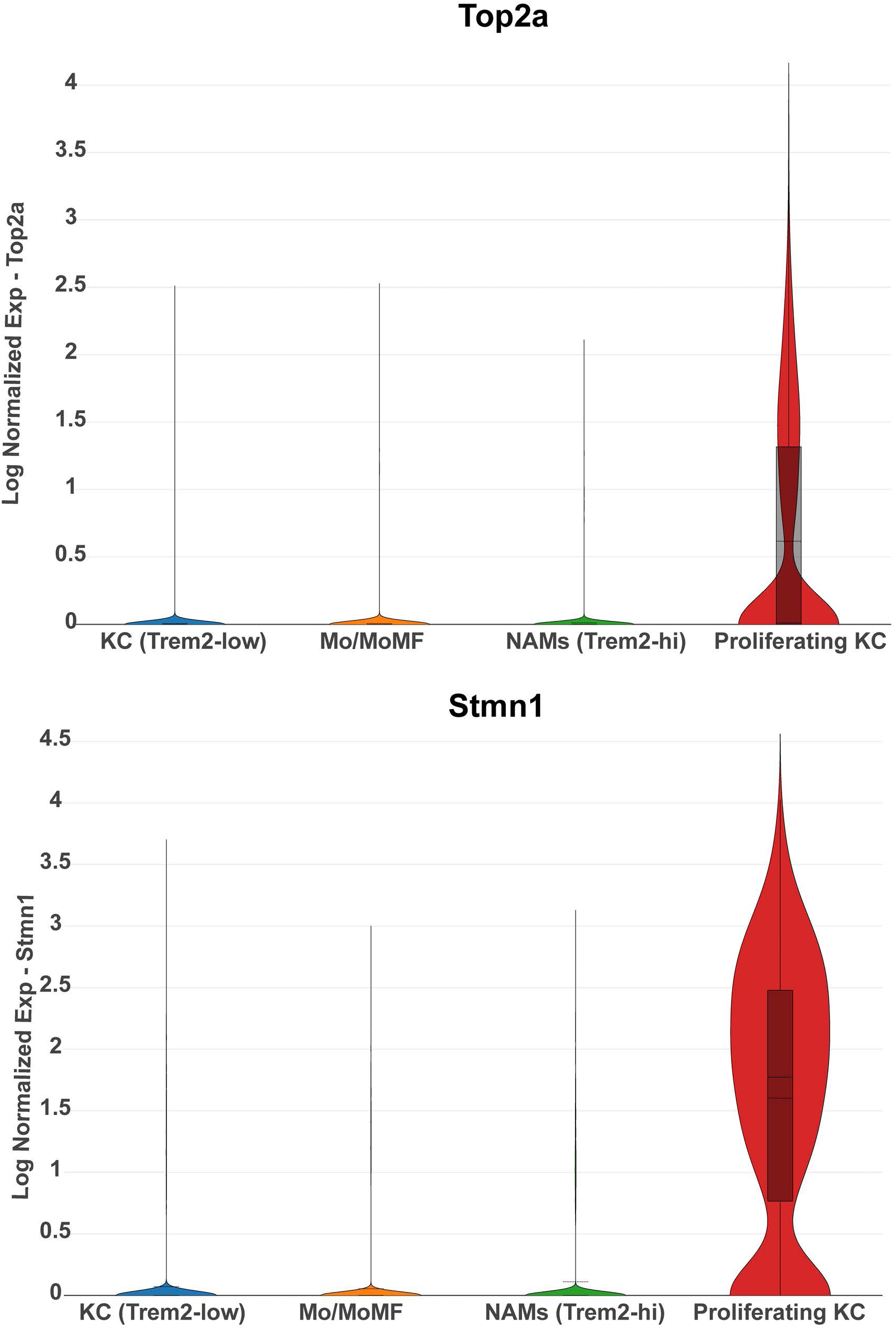
Macrophage proliferation. Violin plots showing expression of genes related to cell division (Top2a and Stmn1) in proliferating Kupffer cells (from Fig. 4G).

